# Invasive Californian death caps develop mushrooms unisexually and bisexually

**DOI:** 10.1101/2023.01.30.525609

**Authors:** Yen-Wen Wang, Megan C. McKeon, Holly Elmore, Jaqueline Hess, Jacob Golan, Hunter Gage, William Mao, Lynn Harrow, Susana C. Gonçalves, Christina M. Hull, Anne Pringle

**Affiliations:** Department of Botany, University of Wisconsin-Madison; Madison, WI, USA; Department of Genetics, University of Wisconsin-Madison; Madison, WI, USA; Department of Biomolecular Chemistry, University of Wisconsin-Madison; Madison, WI, USA; Rethink Priorities; San Francisco, CA, USA; Cambrium GmbH; Berlin, Germany; Department of Medical Microbiology & Immunology, University of Wisconsin-Madison; Madison, WI, USA; Centre for Functional Ecology, Department of Life Sciences, University of Coimbra; Coimbra, Portugal; Department of Bacteriology, University of Wisconsin-Madison; Madison, WI, USA

**Author notes:** Correspondence and requests for materials should be addressed to Yen-Wen Wang.

## Abstract

Canonical sexual reproduction among basidiomycete fungi involves the fusion of two haploid individuals of different sexes, resulting in a heterokaryotic mycelial body made up of genetically different nuclei^1^. Using population genomics data, we discovered mushrooms of the deadly invasive *Amanita phalloides* are also homokaryotic, evidence of sexual reproduction by single individuals. In California, genotypes of homokaryotic mushrooms are also found in heterokaryotic mushrooms, implying nuclei of homokaryotic mycelia also promote outcrossing. We discovered death cap mating is controlled by a single mating-type locus (*A. phalloides* is bipolar), but the development of homokaryotic mushrooms appears to bypass mating-type gene control. Ultimately, sporulation is enabled by nuclei able to reproduce alone as well as with others, and nuclei competent for both unisexuality and bisexuality have persisted in invaded habitats for at least 17 but potentially as long as 30 years. The diverse reproductive strategies of invasive death caps are likely facilitating its rapid spread, revealing a profound similarity between plant, animal and fungal invasions^2,3^.

## Main

Invasion biology focuses on plants and animals and their diseases, while the changing geographical distributions of other microbes go largely unnoticed^4^. The poisonous, European *Amanita phalloides* (the death cap) is an ectomycorrhizal agaricomycete fungus introduced to North America^5,6^. Death caps are now abundant along the California coast and each year cause human and animal fatalities^7,8^. The mechanisms driving the spread of the fungus are not understood. In plants and animals, successful spread is associated with possession of multiple reproductive strategies: After an introduction, the ability to propagate vegetatively and sexually, and to sexually reproduce without a mate, are advantageous^2,3^. We sought to understand how death caps reproduce, and whether selfing is a strategy used by *A. phalloides* to sporulate and move across landscapes.

Agaricomycete fungi are characterized by bisexual reproduction^9^: To complete the life cycle, two haploid mycelia of different sexes (mating types) fuse and form a functionally diploid (heterokaryotic) mycelium. Mushrooms (sporocarps) develop from a heterokaryotic mycelium, undergoing meiosis to produce sexual basidiospores^1^. Like the mycelium, mushrooms carry two different copies of a genome and are heterozygotic. By contrast, unisexual fungi reproduce in the absence of a mating partner^10,11^, and the resulting mushrooms are homokaryotic and without nuclear heterozygosity^1^. In the laboratory, bisexual agaricomycetes can be forced to develop sporocarps from a haploid mycelium with a single mating type in response to physical, chemical, or genetic manipulations^12,13^. However, laboratory mushrooms are typically abnormal^13,14^.

Natural populations with the capacity to generate functional sporocarps both bisexually and unisexually are unknown.

### Invasive, homokaryotic mushrooms

To elucidate the reproductive strategies used by invasive *A. phalloides*, we sequenced genomes of 86 mushrooms collected from Point Reyes National Seashore (PRNS) in California between 1993 and 2015 (N=67), from three sites in Portugal in 2015 (N=11), and from other European countries between 1978 and 2006 (N=8) (Supplementary Table 1). As a fungus grows in a habitat, a single mycelium can develop one or multiple sporocarps, and the 86 mushrooms resolve into 37 distinct genetic individuals (Extended Data Fig. 1a and Supplementary Table 1). Unexpectedly, the estimated heterozygosities of two Californian individuals, g21 and g22, were ten times lower than heterozygosities of other individuals (Fig. 1a). Individual g21 was collected as two mushrooms in 2014 and g22 as six mushrooms in 2004 and 2014. We hypothesized these two Californian individuals are homokaryotic (see also Supplementary Discussion). To test the hypothesis, we used raw read data to identify unique, short DNA strings (unique k-mers) and quantified the number of appearances of each unique k-mer (each k-mer’s depth). Heterozygotic genomes normally show two peaks in k-mer depth, a primary peak and a secondary peak with half the depth of the primary peak (Fig. 1b). The secondary peak is generated by the heterozygous SNPs within a genome and was absent for our putatively homokaryotic individuals (Fig. 1b). In parallel, we investigated sequencing frequencies of alleles at putatively heterozygous sites. Because a true heterozygous site is made up of two alleles, the frequency for each heterozygous allele in sequencing reads should center at 50%, with deviations caused by stochasticity. Once again, putatively homokaryotic individuals are different; the frequency spectra of g21 and g22 are flat (Fig. 1c).

**Fig. 1.**
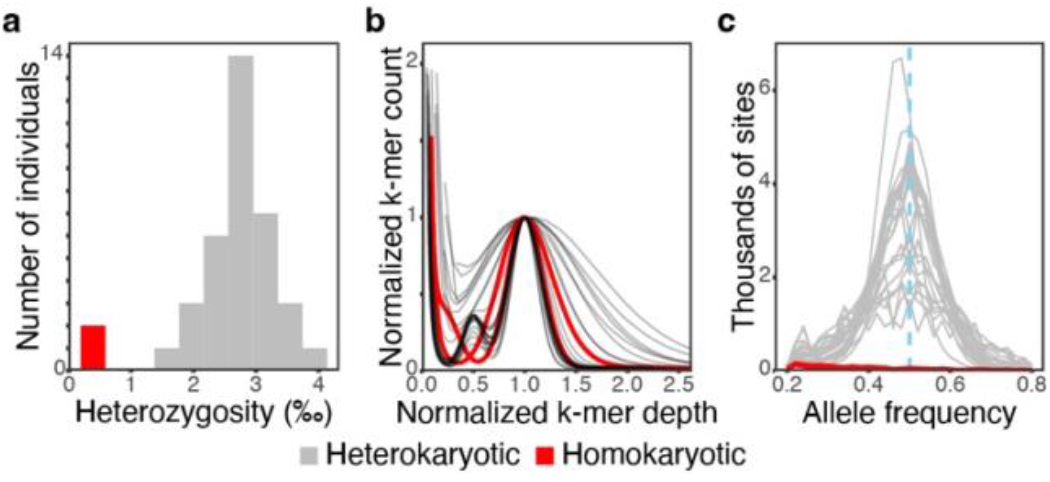
Genomes of two putatively homokaryotic individuals collected in California bear signatures of a haploid or homozygotic genome. (a) Whole-genome heterozygosities of 37 individuals; note the cluster of two Californian individuals at left (in red). (b) Peaks of k-mer depths for Californian and Portuguese individuals; a secondary peak at 0.5 implies heterozygosity and is lacking for the two Californian individuals (in red). (c) Sequencing frequencies of variable SNPs within individuals; peaks at 0.5 indicate heterozygosity and are lacking for the two putative homokaryons.

To determine if homokaryotic individuals can mate with other *A. phalloides* or are reproductively isolated, we estimated kinship among the homokaryotic and other heterokaryotic individuals using a novel algorithm developed by us for use in organisms with mixed haploid/diploid genetic systems, enabling us to identify heterokaryotic individuals housing nuclei from a homokaryon^15^. Many heterokaryotic individuals house either the g21 or g22 nucleus. We cannot distinguish whether these individuals are the parents or offsprings of the homokaryons, nonetheless, because we identified more than one heterokaryotic individual as either the parent or offspring of both g21 and g22, and because only one individual can function as a parent of a homokaryon, we can identify other individuals as offsprings. Homokaryotic individuals appear to mate with other individuals (Fig. 2 and Extended Data Fig. 1b,c). To test whether homokaryotic individuals represent diverged lineages (e.g., cryptic species), we built gene trees from the genomes using 3,324 universal single-copy orthologs (BUSCOs) and combined the gene trees using coalescent-based methods. In the combined tree, homokaryons were neither outgroups nor diverged lineages of the heterokaryotic *A. phalloides* (Extended Data Fig. 2); homokaryotic individuals are not reproductively isolated.

**Fig. 2.**
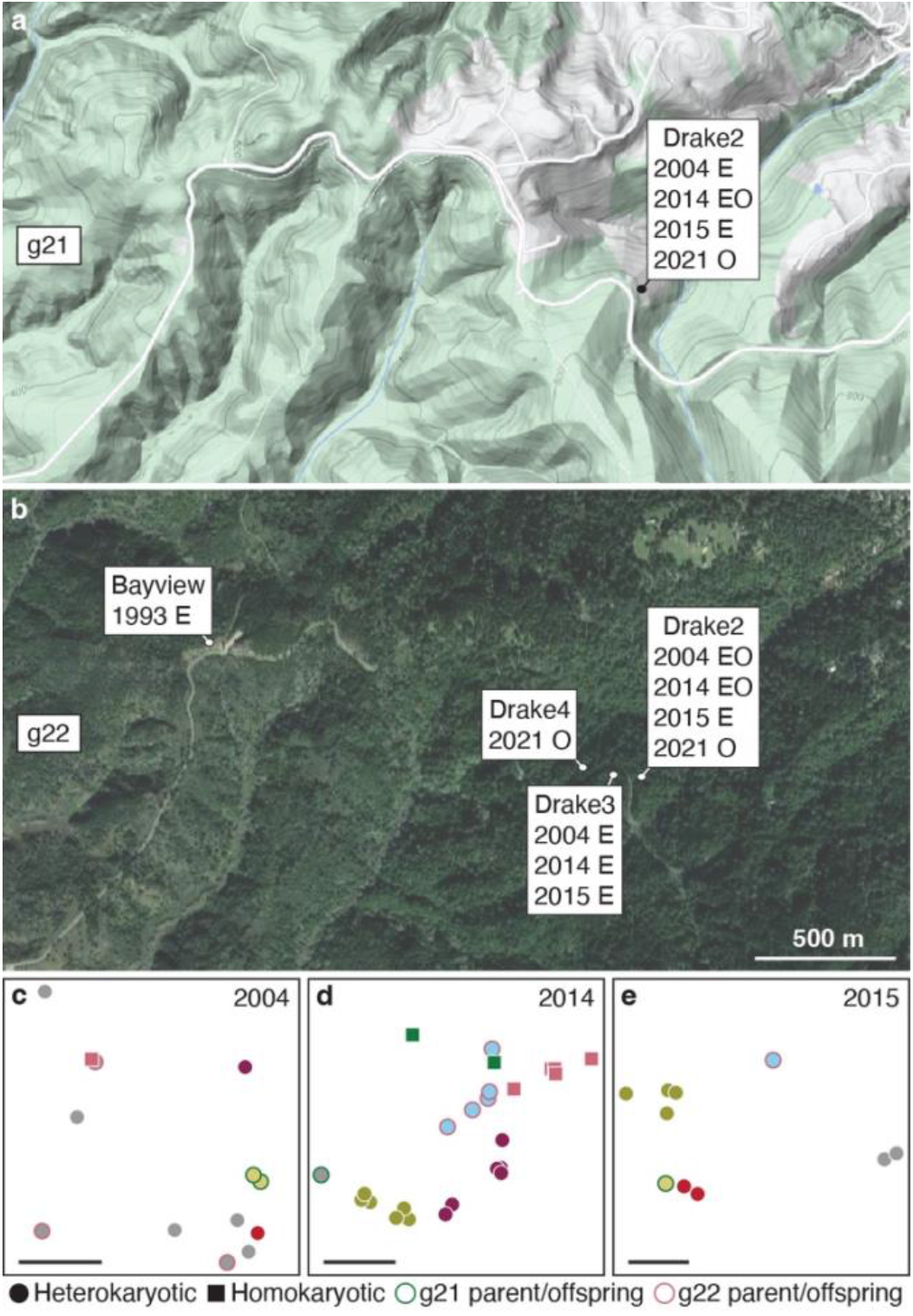
Homokaryotic individuals at Point Reyes National Seashore, California, USA. (a–b) Sites and years from which individuals g21 and g22 were collected; (a) Topography map and g21; (b) Satellite image and g22 (Google, ©2022). O: Site of homokaryotic mushrooms, E: Site of heterokaryotic mushrooms housing the g21 or g22 nucleus, either the parent or offsprings of g21 or g22. (c–e) Fine-scale map of every mushroom collected from Drake2 between 2004–2015. Mushrooms are represented by circles (heterokaryotic mushrooms) or squares (homokaryotic mushrooms of g21 or g22) and shapes of the same color are genetically identical. Gray marks genetic individuals consisting of only a single mushroom. Heterokaryotic parent or offsprings of g21 or g22 are outlined green or rose pink. Scale bars: 5 m.

### Unusual spore numbers

While laboratory sporocarps generated from cultured haploid mycelia are typically aberrant^13,14^, the homokaryotic sporocarps we collected in nature were not very different from the heterokaryotic sporocarps collected from the same sites. In 2021 we revisited PRNS and again collected homokaryotic sporocarps generated by individuals g21 and g22, this time confirming genetic identity by Sanger sequencing of 11 loci (Supplementary Tables 2 and 3). Collected sporocarps are morphologically similar to heterokaryotic sporocarps (Fig. 3a,b). Using dried materials from the original collections made in 2014, we discovered that both homokaryotic and heterokaryotic sporocarps possess unisporic, bisporic, and trisporic basidia, as well as canonical tetrasporic basidia (Fig. 3c,d and Extended Data Fig. 3). However, the ratios of the spore arrangements were different among the five individuals we measured (Fig. 3d), with heterokaryotic g25 possessing the highest frequency of tetrasporic basidia and homokaryotic g21 possessing the highest frequency of unisporic basidia. Next, we imaged patterns of nuclei within basidia and basidiospores. As documented in closely related *Amanita* species^16^, younger basidiospores house one nucleus, and more mature spores house two, likely the result of a mitotic division within developing basidiospores (Extended Data Fig. 4a,b). Tetrasporic basidia leave no nuclei behind in the originating basidium (Extended Data Fig. 4a,b), but in trisporic basidia, one nucleus remains in the basidium (Fig. 3e,f and Extended Data Fig. 4c). The number of spores on a basidium does not appear to influence nuclear segregation; regardless of spore number, each spore receives one meiotic nucleus from the originating basidium, a phenomenon also observed in bisporic *A. bisporigera*^17^. Pseudohomothallism, or the phenomenon of a spore with two genetically different nuclei, and hence, the ability to grow and sexually reproduce without a mate^10^, is not a feature of the *A. phalloides* life cycle. The same spore-nuclear dynamics are observed in both heterokaryotic and homokaryotic sporocarps, suggesting basidia of both kinds of sporocarps are cytologically similar. Moreover, fluorescent staining of nuclei in mycelia taken from stipe tissues revealed cells of both homokaryotic and heterokaryotic sporocarps can be multinucleate (Extended Data Fig. 4d,e).

**Fig. 3.**
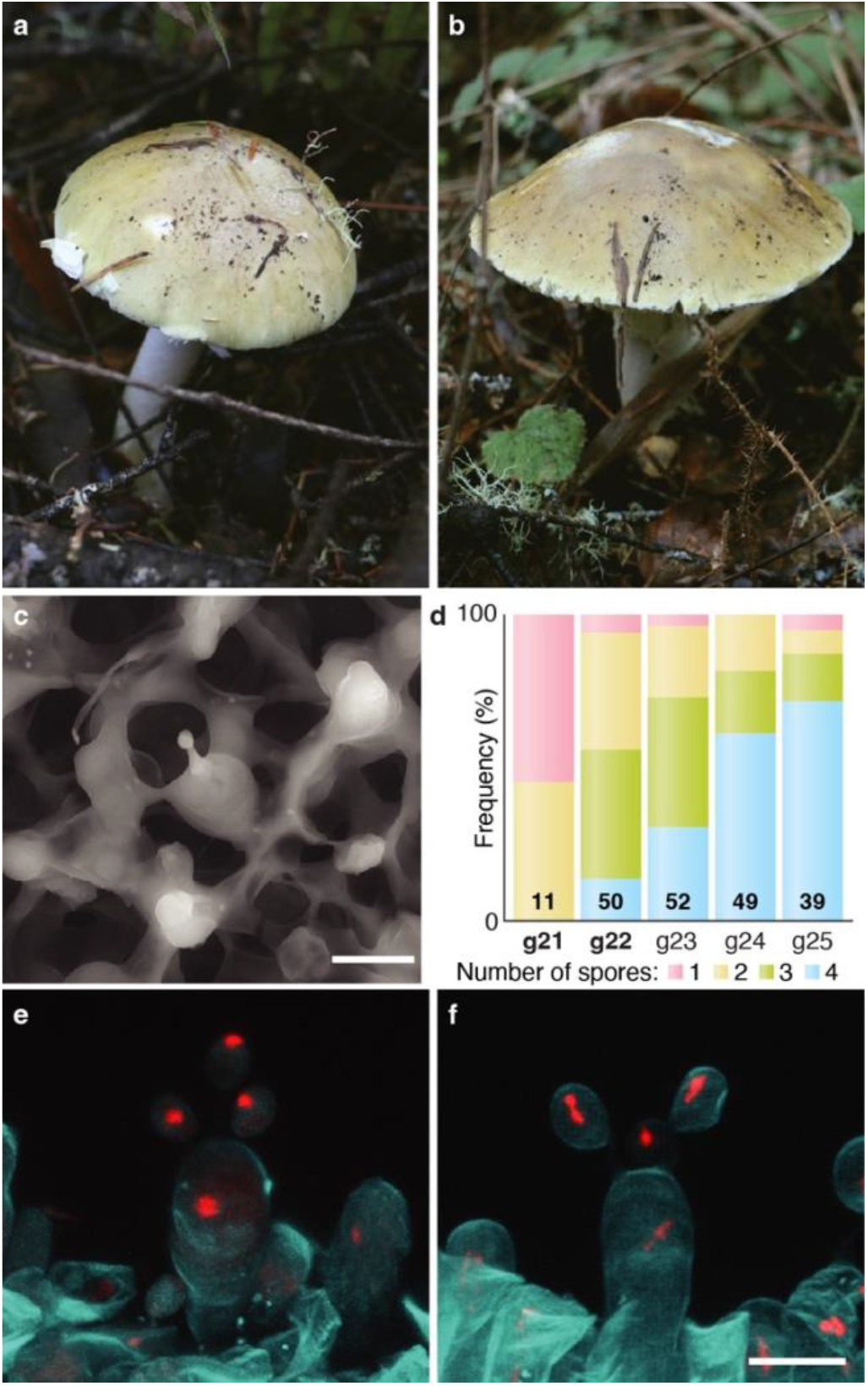
Morphology of heterokaryotic and homokaryotic sporocarps. (a) Heterokaryotic sporocarp found in 2021. (b) Homokaryotic sporocarp found in 2021. (c) Scanning electron microscopy of a unisporic basidium from a homokaryotic sporocarp. (d) Frequency spectra of the number of spores per basidium among five individuals. Homokaryotic individuals are bold (g21 and g22). Number of basidia counted indicated within each column. (e–f) Z-stack composite image of confocal microscopy of trisporic basidia from heterokaryotic (e) and homokaryotic (f) sporocarps; in (f) the basidium was more mature and nuclei are dividing. Red: Vybrant Orange (nuclei); cyan: Calcofluor White (cell wall). Scale bars: 10 μm.

### Genetics of sexual reproduction

Because sporocarp development is closely associated with sexual reproduction, we next sought to determine, for the first time, the genetics of a mating system within the genus *Amanita*. Canonical agaricomycete mating systems are controlled by two mating (MAT) loci: a pheromone and pheromone receptor locus (P/PR) and a homeodomain locus (HD). Successful sexual reproduction requires two fusing, haploid mycelia to carry different alleles at P/PR and HD, a system termed tetrapolar heterothallism. In bipolar heterothallic systems, either the P/PR and HD are linked, or one locus (usually P/PR) is no longer involved in mating^9^.

We discovered *A. phalloides* possesses a bipolar mating system (Supplementary Discussion). As anticipated, we identified two putative pheromone receptor (*PR*) genes (Fig. 4a and Extended Data Figs. 5a and 6); however, we found that among *Amanita* species the number of putative *PR* genes is not consistent (ranging from two to five), and *PR* genes reside in a genomic region only weakly syntenic across the genus (Extended Data Fig. 5a). Unexpectedly, we were unable to identify genes predicted to encode pheromones (*P* genes) near the *PR* genes in *A. phalloides*. Instead, apparent homologs of pheromones are located in other regions of the genome (Supplementary Fig. 1). MAT loci are typically highly diverse, the result of frequency-dependent selection^18–20^, but in *A. phalloides*, putative *PR* genes exhibit low genetic diversity (Fig. 4a). Moreover, nine of the 25 heterokaryotic individuals tested carry identical copies of the *PR* genes. Finally, the two *PR* genes are orthologous to non-mating type-determining genes in other fungal species, as demonstrated in a species-tree-aware gene phylogeny (Extended Data Fig. 7). The apparent absence of pheromones, as well as low genetic diversity, functional homozygosity of *PRs* in heterokaryotic individuals, and orthology between *PRs* and non-mating type-determining genes in other fungi, each suggests the *PR* genes are not involved in mating.

**Fig. 4.**
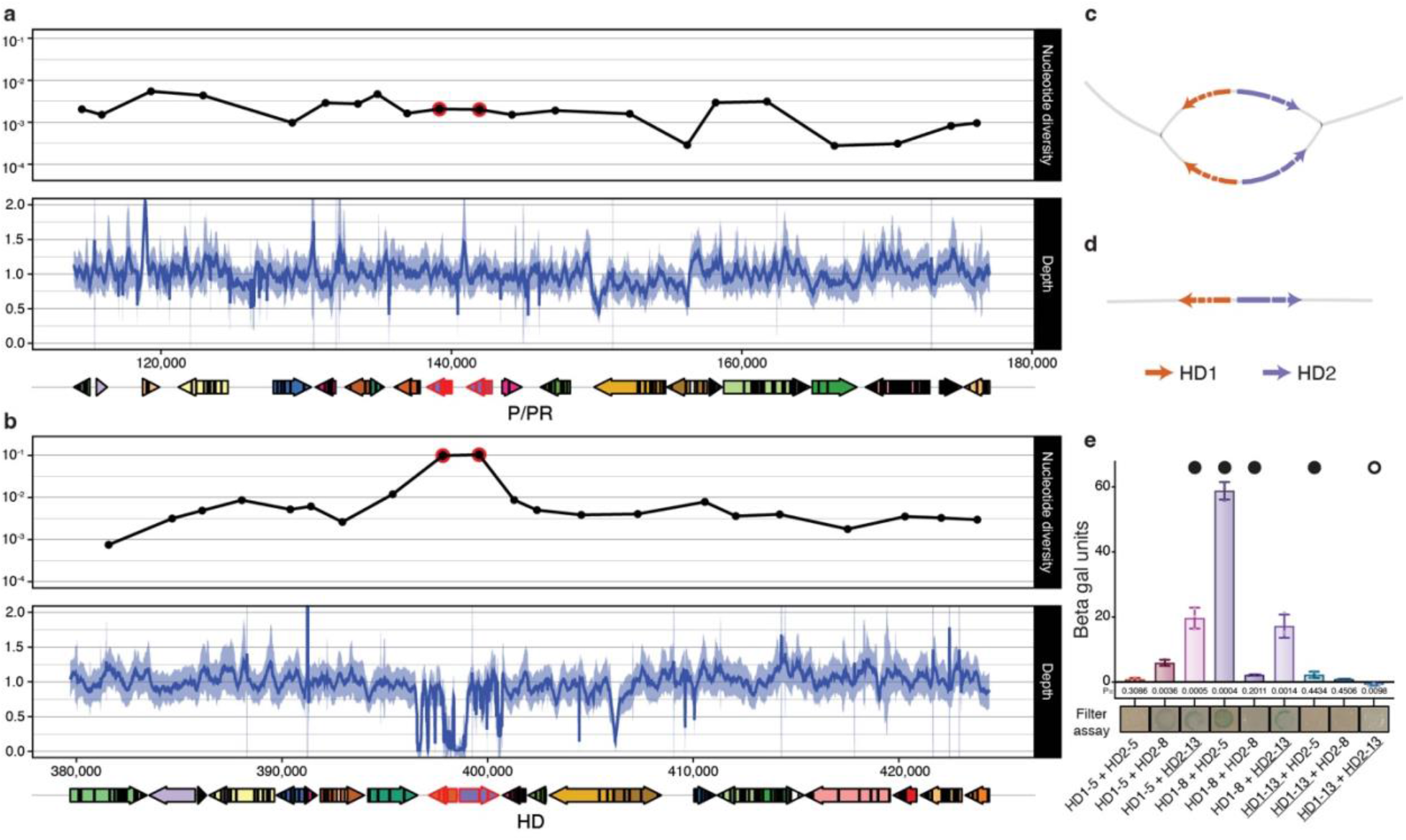
Genomic features of putative mating loci and evidence for interactions between *HD* genes. (a–b) Nucleotide diversity, sequencing depth and gene orientation of P/PR (a) and HD (b) loci. *PR* and *HD* genes outlined in red. Nucleotide diversity calculated from *de novo* genome assemblies; note diversity increase and depth drop at HD locus. (c–d) Assembly string graphs of the HD locus of heterokaryotic (c) and homokaryotic (d) sporocarps. Homokaryotic genomes house one copy of the locus. (e) Yeast two-hybrid β-galactosidase reporter activities from strains harboring HD1 and HD2 proteins of different alleles (designated 5, 8, and 13; HD1s fused with a Gal4 activation domain and HD2s fused with a Gal4 DNA binding domain; allele 13 (underlined) identified in a homokaryon (g22)). Closed circles: HD1 and HD2 combinations found in heterokaryotic mushrooms; Open circle: HD1 and HD2 combination found in homokaryotic mushroom. Error bars: standard errors of the means (each includes three biological replicates). P-values are determined by comparing *LacZ* expression in β gal units (Miller Units) to each respective BD plasmid co-transformed with the empty AD plasmid in an unpaired two-sided Student’s t-test (d.f. = 4)

In contrast to the *PR* genes, *HD* genes in *A. phalloides* appear typical of MAT loci in agaricomycete fungi. In a typical MAT locus, there are two *HD* genes, designated as *HD1* and *HD2*. If the alleles of *HD1* and *HD2* in the nuclei of a heterokaryon encode compatible proteins, the proteins can form heterodimers, and heterodimers function as transcriptional regulators to promote mating and sexual development^21^. Across the genus *Amanita*, the HD locus also consists of two *HD* genes (Fig. 4b and Extended Data Fig. 5b). Within *A. phalloides*, the numbers of raw reads mapped onto the reference genome at the two *HD* genes were very low, compared to adjacent genes (Fig. 4b), suggesting HD alleles are highly diverged from each other. To extract the precise sequence of the alleles of each gene in each individual, we *de novo* assembled a genome for each Californian and Portuguese sporocarp (N=77) (Fig. 4c,d and Extended Data Fig. 8). In all assembled genomes, we observed nucleotide diversity peaking at the *HD* genes, a hallmark of diversifying (frequency-dependent) selection (Fig. 4b). As expected, both the HD1 and HD2 proteins were predicted to possess canonical homeodomains (Extended Data Fig. 9), and only the *HD1* genes encode nuclear localizing signal (NLS) peptides. Models of predicted heterodimers suggest that HD1 and HD2 interact with one another via their N-termini (Extended Data Fig. 9). The pattern demonstrates a congruence between *A. phalloides HD* genetic signatures and the known functionality of *HD* genes in *Coprinopsis cinerea*^21^; in *Co. cinerea* the NLS on HD1 is required to import HD2 into the nucleus where it can act. In the aggregate, our data document *A. phalloides* as a bipolar species in which mating types are determined by a single MAT locus containing only *HD* genes.

Four mechanisms can explain the development of a sporocarp from a haploid fungus^1^ (but see also Supplementary Discussion^22^): (1) gene conversion from a silent mating type locus to create compatible mating loci, (2) gene duplication of the mating type locus followed by divergence to generate compatible mating loci within a single genome, (3) self-compatibility of genes within a mating type locus, or (4) drivers enabling a bypass of mating type control^1^.

Neither gene conversion nor gene duplication can explain the homokaryotic *A. phalloides* sporocarps because we did not find multiple copies of either *HD* gene within any homokaryotic genome. To test for self-compatibility within the *A. phalloides* MAT locus, we carried out protein-protein interaction tests between the HD1 and HD2 proteins. We hypothesized that if the HD1 and HD2 proteins from the homokaryotic sporocarp are self-compatible, they would interact with one another in a yeast two-hybrid assay (Supplementary Tables 4 and 5). We evaluated HD1 and HD2 proteins from three different MAT alleles; two from heterokaryotic samples (alleles designated 5 and 8), and one from a Californian homokaryotic sample (allele 13). For the heterokaryotic HD1 and HD2 alleles, we anticipated that at least one of the HD1-HD2 pairs from different MAT alleles would interact and that the HD1 and HD2 proteins from the same MAT allele would not interact. This was, indeed, the case and is congruent with findings in other fungal systems (Fig. 4e and Extended Data Fig. 10)^21,23,24^. The HD1 and HD2 proteins from the homokaryotic sporocarp also did not interact with each other or with themselves under the conditions tested (Fig. 4e and Supplementary Fig. 2). The heterodimerization of HD proteins from the homokaryotic MAT allele is unlikely to enable sporocarp development. Our finding suggests other mechanisms besides heterodimerization enable the development of homokaryotic mushrooms.

### Geography of unisexual individuals

To discover whether homokaryotic *A. phalloides* sporocarps can be found at other sites or in other ranges, we Sanger sequenced the variable beta-flanking gene adjacent to *HD1* as well as ten other mostly unlinked conserved genes from an additional 109 sporocarps collected from three sites in and around Berkeley, California (N=15), from populations introduced to New Jersey (N=15) and New York (N=10), and from Canada, where introduced death caps grow on Victoria Island (N=8). From its native range, Europe, we sequenced a population of death caps collected near Montpellier, France (N=12), from two sites in Norway (N=6), twelve sites in the UK (N=14), four sites in Austria (N=13), two sites in Estonia (N=2), two sites in Hungary (N=12), and one site in Switzerland (N=2) (Supplementary Table 6). The sequence data allow us to evaluate whether collected sporocarps lack heterozygosity. With this approach the probability of a heterokaryotic sporocarp being misidentified as homokaryotic is estimated as lower than 0.2%. We discovered no homokaryotic sporocarps at any other site (Supplementary Table 7).

Homokaryotic sporocarps appear to be extremely rare in nature, but in California homokaryotic individuals can span large territories, and sporocarps of g22 were found up to 200 m apart (Fig. 2b). These homokaryotic individuals have persisted for at least seven (2014–2021; g21) and up to 17 (2004–2021; g22) years. An herbarium specimen collected in 1993 at a site approximately 1.6 km away from our first g22 collection is heterokaryotic and is either the parent or offspring of individual g22. If it is an offspring, the g22 nucleus would have persisted for nearly 30 years in California and would have a much wider territory than we have discovered using only the presence of its homokaryotic mushrooms as a guide. Its apparent independence recalls nuclear dynamics in other fungi^25^.

### The spread of invasive mushrooms

The spread of *A. phalloides* in California is likely facilitated by its ability to sporulate without mating with another individual. The fungus is both unisexual and bisexual, revealing a previously unsuspected reproductive flexibility in a natural population of death caps.

Homokaryotic mycelia are considered an ephemeral stage of agaricomycete life cycles^26^. Our discovery of nuclei able to live alone (as homokaryons) as well as with other nuclei (in heterokaryotic mycelia) is surprising, as is the persistence of these nuclei at PRNS for at least 17 (and up to 30) years. We hypothesize single basidiospores occasionally germinate into haploid mycelia able to develop mushrooms and sporulate *via* endoduplication. Some of the offspring of these mushrooms mate, while others do not, and the cycle repeats. Alternatives to canonical heterothallic mating in fungi are typically associated with pseudohomothallism or mating type switching^1^, but neither mechanism explains the homokaryotic death cap sporocarps. *A. phalloides* unisexuality appears most similar to the unisexual reproduction described in *Cryptococcus* species, in which mating type control is bypassed to develop basidia and spores. Unisexuality in both *Cr. gattii* and *A. phalloides* is associated with biological introductions^27^, providing additional support for the selective advantage provided by self-reproduction in an introduced range, and revealing a profound similarity between plant, animal, and fungal invasions.

## Supporting information

Extended Data, will be used for the link to the file on the preprint site.

Supplementary Information, will be used for the link to the file on the preprint site.

## Acknowledgments

We thank Benjamin Becker, Tom Horton, Benjamin Wolfe, Cat Adams, Tom Bruns and the Bruns laboratory, Sydney Glassman, Brenda Callan, Lynne Boddy, the Pringle laboratory, Franck Richard, Milton Drott, Debbie Viess, Forrest Gander, and Ashwini Bhat for help collecting mushrooms, and fungaria for providing us with additional samples from stored collections. Field work at PRNS was conducted under permits granted to the Bruns laboratory and Y. W. (PORE-2021-SCI-0047). For library preparation, we used the University of Wisconsin – Madison Biotechnology Center’s DNA Sequencing Facility (Research Resource Identifier – RRID:SCR_017759) for genomic DNA and we used Biotechnology Center’s Gene Expression Center Core Facility (RRID:SCR_017757) to prepare the RNA library. To sequence, we used the DNA Sequencing Facility (RRID:SCR_017759) for both genomes and transcriptome. We acknowledge various grants and fundings enabling this project, including University of Wisconsin-Madison (A.P., Y.W.), Human Frontier Science Program grant RGP0053 (A.P.), Fulbright U.S. Scholar grant (A.P.), Mycological Society of America Graduate fellowship (Y.W.), National Institutes of Health grant T32 GM007133 (M.C.M.), Fundação para a Ciência e Tecnologia COMPETE Programme and National Funds FEDER fund PTDC/BIA-BIC/122142/2010 (S.C.G.), National Institutes of Health grant R01 AI137409 (C.M.H.).

## Author contributions

Y.W., M.C.M., H.E., J.H., J.G., H.G., C.M.H. and A.P. designed experiments. A.P., Y.W., J.G., H.E. and S.C.G. led field work and Y.W., M.C.M. and H.G. performed laboratory experiments. Y.W., M.C.M., H.E., J.H., J.G., W.M., L.H. and A.P. analyzed data. A.P. and C.M.H. supervised the study. A.P. obtained funding. Y.W. wrote the initial manuscript. All authors reviewed and provided feedback towards the final manuscript.

## Foot notes

Supplementary Information is available for this paper.

Correspondence and requests for materials should be addressed to Yen-Wen Wang.

## Methods

### Mushroom collecting, genome sequencing, annotation, and variant calling

Sporocarps were collected from various herbaria and during three expeditions to Point Reyes National Seashore (PRNS), California in 2004, 2014 and 2015, and in 2015 from three sites in Portugal. A total of 86 sporocarps were collected: 67 Californian sporocarps (one early herbarium sample dates to 1993), 11 Portuguese sporocarps, and eight sporocarps from other European countries (Supplementary Table 1). Specimens of sporocarps are deposited in the fungarium in Pringle laboratory. The Californian specimens collected from 2004 to 2015 were mapped.

Details of genome sequencing, annotation and variant calling are described in Supplementary Methods. In brief: A reference genome (mushroom 10511) was sequenced with both Illumina and PacBio platforms and assembled using a hybrid approach. A transcriptome (mushroom 10721) was sequenced using an Illumina platform to enable genome annotation with the GenSAS web server^28^. Illumina genome sequences of other mushrooms were mapped to the reference genome and single nucleotide polymorphisms (SNPs) called using the Genome Analysis ToolKit (GATK)^29^. SNPs were filtered based on sequencing depth, presence of transposable elements, and putative contamination.

### Heterozygosity estimation, k-mer analysis and allele frequencies

We tested whether any sporocarps without heterozygosity are found among our sequenced individuals by comparing individuals’ heterozygosities, k-mer distributions and allele frequency graphs. Because a single mycelium can generate multiple mushrooms, we first sorted the sporocarps into genetic individuals (see Supplementary Methods), and then we counted the homozygous sites for each sporocarp using the filtered variant calling format (VCF) file and VCFtools ver. 0.1.16^30^. The heterozygosity of each sporocarp was next estimated with

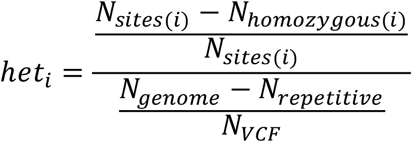

where *N*_*sites(i)*_ was the number of non-missing sites in sample *i. N*_*homozygous(i)*_ was the number of homozygous sites in sample *i. N*_*genome*_, *N*_*repetitive*_ and *N*_*VCF*_ were the assembly size, the size of the masked repetitive regions identified by REPET, and the number of sites in the VCF file. The heterozygosity of each genetic individual was calculated as the mean of the heterozygosities of all sporocarps making up that individual.

To further explore the zygosities of individuals with low estimated heterozygosities, we first used k-mer analyses. Because herbarium sporocarps (most non-Portuguese European sporocarps and the 1993 Californian sporocarp) had lower DNA quality than the newer specimens collected by us, we only compared our samples from California or Portugal collected after 2000. For k-mer analysis, we used BBMap ver. 38.73^31^ to generate k-mers of 23 bp and count the number of each different k-mer. The k-mer frequency distribution of each genome was normalized by its largest peak.

As a second approach to probe zygosities, we investigated per individual allele frequency. We plotted histograms of the sequencing frequencies of alleles of putatively heterozygotic sites from the filtered VCF. Heterozygotic diploid individuals should display a peak at 0.5 because the two alleles at a site normally have a similar sequencing depth. To be sure sequencing depth was unaffected by the variant re-calling step, we also reran this analysis using the same filters used to generate the filtered VCF but without variant re-calling.

### Kinship analysis

To test if the homokaryotic individuals discovered in California are mating with other individuals in the population, we estimated kinships between homokaryotic individuals and heterokaryotic individuals with KIMGENS, a population-structure-robust estimator^15^. We also used KIMGENS to confirm the results of a clone correction based on Euclidean genetic distances (Supplementary Methods). To generate a set of SNPs for KIMGENS, we first removed sites called as heterozygous in any homokaryotic sporocarp from the filtered VCF file. Following KIMGENS, we set a threshold at 0.75 to determine which mushrooms belong to the same homokaryotic individual(s) and at 0.4375 to determine which mushrooms belong to the same heterokaryotic individual(s). The results from KIMGENS and Euclidean distances were identical.

In subsequent analyses we generated a consensus genotype for each genetic individual consisting of multiple mushrooms. We next compared kinships among the 37 genetic individuals. Because kinship is defined as the probability of IBD between two randomly chosen alleles, one from each individual in a pair of interest, the immediate kins of a homokaryotic individual (i.e. its parent and heterokaryotic offsprings) will share kinships of 0.5 with the homokaryotic individual.

### Population-level phylogeny reconstruction

Although kinship estimates suggest homokaryotic individuals are mating, they cannot exclude the possibility of homokaryotic individuals belonging to other (as yet unidentified and unnamed) reproductively isolated groups (e.g., a cryptic species). To explore this hypothesis, we used a coalescent-based phylogeny reconstruction method. We called BUSCOs of Agaricales from the reference assemblies of *A. phalloides* and *A. subjunquillea* (the closest relative of *A. phalloides*) with BUSCO ver. 5.2.2 using *Laccaria bicolor* for the AUGUSTUS species parameter. Then, the DNA sequences of the common single-copy BUSCOs of each individual were called from filtered VCF files with *vcf-consensus* from VCFtools with IUPAC codes. All sequences for a BUSCO were aligned with MAFFT ver. 7.427^32^, and a phylogeny was reconstructed using IQ-TREE ver. 2.0.6^33^ with a nuclear substitution model identified by ModelFinder Plus^34^ and bootstrapped 1000 times with a hill-climbing nearest neighbor interchange (NNI) search^35^. Then, we controlled for highly related individuals in the phylogenies by using a Markov clustering analysis (MCL). To avoid excess pruning, any pairwise kinship lower than 0.1 was recoded to 0 and the inflation rate of MCL was set to 1.5. To ensure one homokaryotic individual was included in each of the final phylogenies, one individual was picked for each cluster manually. Since the two homokaryotic individuals were related, this pruning process retained only one homokaryotic individual: g22. We summarized the phylogenies of BUSCO genes using the “constrained-search” branch of ASTRAL ver. 5.6.9^36,37^. We reconstructed two phylogenies: one unconstrained, and the other constrained by forcing *A. subjunquillea* and the homokaryotic individual to form a single cluster.

### Identification of mating type loci across the genus *Amanita*

To establish if the two homokaryotic individuals possess a single mating type allele, we first identified mating type loci in the reference genome and across the genus *Amanita*. Basidiomycetes are typically tetrapolar and so we used the protein sequences of the mating-type homeodomain proteins (HD; XP_001829154.1, XP_001829153.1) and pheromone receptors (PR; AAQ96344.1, AAQ96345.1) of *Coprinopsis cinerea* as queries for BLASTp^38^, searching within the predicted proteome of *A. phalloides* (JAENRT000000000) to identify homologs. To identify syntenic regions, we searched for the *HD* and *PR* genes, as well as homologs of ten genes located up- and downstream of the two loci, in the predicted proteomes of *A. brunnescens* (JNHV00000000.2), *A. polypyramis* (JNHY00000000.2), *A. muscaria* var. *guessowii* (JMDV00000000.1) and *A. inopinata* (JNHW00000000.2). After completing structural analyses of proteins (see below), we manually reannotated the genes of the two loci.

### Identification of pheromones in *A. phalloides*

The genes of pheromones (*P* genes) are shorter than canonical genes and our annotation pipeline was unable to identify pheromones within the pheromone and pheromone receptor (P/PR) locus. To identify pheromones, we first used the ORFfinder^39^ in NCBI to predict open reading frames (ORFs) longer than 30 bp between 5 kbp upstream and downstream of the *PR* genes. Then we looked for any ORFs with ER/DR (N-terminal cleavage) and -CaaX (farnesylation) motifs^21^. However, no pheromones close to the *PR* genes were identified, and so we used protein sequences of the pheromones of *A. muscaria* (which appears to have retained its *P* genes at the P/PR locus) taken from the JGI genome portal (gene ID: 163418 and 163420) to search in the *A. phalloides* genome assembly with tBLASTn. These pheromone gene predictions were further refined by comparing the predictions against transcripts.

### Sequencing depth of mating type loci

Fungal mating type locus alleles are in general highly polymorphic, and read mapping algorithms often perform poorly on them, resulting in erroneous variant calls. To understand if poor mapping is a major concern for the mating type loci of *A. phalloides*, we used SAMtools ver. 1.5^40^ to estimate the sequencing depth of each locus and normalized each locus’s average depth to one.

### Identification of homeodomains proteins in *A. phalloides* I: *De novo* assembly for HD calling among all samples

We discovered the *A. phalloides* HD locus does in fact map poorly, and so we chose to *de novo* assemble the genomes of each sporocarp as the first step towards identifying the HD locus of every genetic individual. In a graph-based genome assembly pipeline, different alleles of a locus in a heterokaryotic genome will be assembled into two unitigs (equivalent to a contig without conflicting nucleotides). Because these two unitigs represent two alleles, they are also called haplotigs. In a successful assembly, the two haplotigs each connect with the rest of the genome on either side, and so form a “bubble” within the assembled genome. We took advantage of assembly bubbles to identify the *HD* genes.

We first trimmed the raw reads with Trimmomatic ver. 0.35^41^ to remove the adapters tagged as “ILLUMINACLIP:TruSeq3-PE-2.fa:2:30:10” and “MINLEN:75”. We then assembled each sample with the de Bruijn graph-based assembler SPADES ver. 3.13.1^42^, using both paired-end reads and unpaired reads (k-mer size=21). To identify the unitigs containing *HD* genes (the HD haplotigs), we used the DNA sequences of *HD* genes from the reference genome of *A. phalloides* as queries to search within the assembly graph with Bandage ver. 0.8.1^43^. To confirm the HD haplotigs as the two alleles of a sample, we also identified the connected unitigs five unitigs away/around the HD haplotigs on either side of them. We grouped sporocarps into one of five categories (Extended Data Fig. 8): 1. closed bubble: the HD haplotigs attached to the same unitigs on both sides; 2. open bubble: the HD haplotigs only attached to the same unitig on one side; 3. detached: the HD haplotigs were not linked to each other; 4. complexed: more than one unitig on either side of the HD haplotigs were linked back to the haplotigs; and 5. odd-number unitig: when only one or more than two unitigs were identified as HD haplotigs. For individuals appearing to have only one HD haplotig, we evaluated assembly quality by mapping the raw reads back to the HD haplotig using the BWA MEM algorithm.

### Identification of homeodomains proteins in *A. phalloides* II: Annotation of *HD* genes in each genome

Using the genome assemblies of sporocarps, we annotated the *HD* genes of every specimen collected between 2004 and 2015 from California and Portugal. We first used four different strategies to predict the genes within each genome’s HD haplotigs: a pretrained AUGUSTUS, a self-trained AUGUSTUS, SNAP and CodingQuarry. For pretrained AUGUSTUS, we used the AUGUSTUS web interface^44^ to predict genes based on a *Laccaria bicolor* gene model. For self-trained AUGUSTUS, we used the gene models chosen from whole genome annotation and trained AUGUSTUS ver. 3.3.3 on Galaxy server^45^. For SNAP and CodingQuarry, we followed the strategies used in whole genome annotation described above.

Gene predictors sometimes produce faulty annotations, so next we manually reannotated the introns, and start and stop codons, of the *HD* genes of 10721 (the same specimen of which transcriptome was sequenced) based on its mapped transcriptomic reads. Fungi normally use the first start codon in a transcript as the translation initiation site^46^, and therefore we annotated the first in-frame AUG of the majority of transcripts as the start codon. We noted that 15 of 986 transcriptomic reads extend toward the upstream of most other reads of the *HD2* gene. These 15 reads include another putative start codon starting at the third nucleotide. But because of the low depth of these reads and the close position of the alternative putative start codon to the 5’ end of the transcript, we did not consider these 15 reads as encoding the true start codon of *HD2*. Using this information, we later manually reannotated the *HD* genes for each sporocarp’s genome by aligning and comparing among protein sequences. Because the sporocarps making up a genetic individual did not always have the exact same DNA sequences, we first aligned the protein sequences with MAFFT and reverse translated the alignment to coding DNA sequences (CDSs). Then we generated the consensus CDS for each allele of each individual.

To compare sporocarps to each other, and to discover whether HD alleles are shared across different genetic individuals, we reconstructed phylogenies for the *HD1* and *HD2* alleles. We translated the already aligned CDSs back to proteins and built a phylogeny for each of the two *HD* genes with RAxML-HPC ver. 8.2.9 with the best protein substitution models determined by ModelTest-NG^47^ and bootstrapped for 100 times.

### Structural analyses of pheromone receptor and homeodomain proteins

Next, we sought to understand if the *PR* and *HD* genes encode canonical mating type determining proteins by analyzing protein structures. We first explored whether either of the *PR* genes encodes the canonical structures of the seven-transmembrane domain in G-protein coupled receptors (GPCRs) or signal peptides. We identified the sequences from the reference genome annotated as *PR* genes. To identify transmembrane helices, we submitted sequences to the CCTOP server^48^ with TM filter. To identify signal peptides, we used SignalP 5.0^49^ and searched for eukaryotic signal peptides. Then we predicted the general structures of the *PR* genes using Alphafold2^50^ on Cosmic^2 51^ and its full protein database (February, 2022).

Then, we chose to predict the HD protein structures of heterokaryotic g19 (alleles 8 and 13), g33 (alleles 5 and 8) and homokaryotic g22 (allele 13), and targeted nuclear localization signal (NLS) peptides and homeodomains. To identify the NLS, we used cNLS Mapper^52^ with a cut-off score of 5.0, and we only searched for bi-partite NLS within 60 amino acids of either terminal of the protein. To identify homeodomains, we used RaptorX-Property^53^ to first identify alpha helices using a 3-class classification (of alpha-helix, beta-sheet or coil). Then we searched for alpha helices homologous to the homeodomains^54^. To explore potential interactions between HD proteins, we used Alphafold-Multimer^55^ on Cosmic^2^ with its full protein database to predict the heterdimeric structure of HD1-8 and HD2-5 which were encoded by genes on the different alleles of a single individual: g33 (February, 2022). The protein structure was visualized with PyMol ver. 2.4.1^56^ and the predicted alignment errors (PAEs) were extracted with paem2png.py (https://github.com/CYP152N1/plddt2csv).

### Nucleotide diversity of putative MAT loci

To test if the *PR* and *HD* genes are under diversifying selection, we compared their nucleotide diversities to the diversities of upstream and downstream genes. Because the *HD* genes were identified using a different approach from other genes, we were unable to filter SNP sites using a single protocol. Therefore, we used the unfiltered VCF of only heterokaryotic sporocarps from California or Portugal, and only genomes for which the HD locus was assembled correctly. We used the R package *pegas*^57^ to calculate the nucleotide diversities of the ORFs of *HD* genes. We used VCFtools with --site-pi flag to calculate the nucleotide diversities of other genes and correct for gene length as necessary. In this analysis, we did not clone correct.

### Interspecies phylogeny of *PR* genes

Because no substantial diversity was present within the *PR* genes of our population genomics dataset and many fungal species have *PRs* that do not determine mating type, we investigated the orthology of *A. phalloides* and other model systems’ PR proteins using a species-tree-aware gene phylogeny. We first harvested the PR sequences of species listed in Coelho *et al*. (2017)^9^ from the JGI genome portal^58^ using the gene ontology term for mating-type factor pheromone receptor activity (GO: 0004932), and sequences of two ascomycete pheromone receptors (*Saccharomyces cerevisiae* and *Pneumocystis carinii*) from NCBI (Acc. No. P06783.1 and AAG38536.1). We then used MAFFT to align the protein sequences of the retrieved pheromone receptors and our five *Amanita* species, trimming the alignment with trimAl ver. 1.4.rev15^59^. Finally, we reconstructed a species-tree-aware gene phylogeny with GeneRax ver. 2.0.4^60^ using the best substitution model identified by ModelTest-NG, the species tree as described in Coelho *et al*. (2017)^9^, and an undated duplication-loss model.

### Yeast two-hybrid of *HD* genes

We conducted a yeast two-hybrid experiment to test whether proteins encoded by the *HD1* and *HD2* of different alleles in a heterokaryotic individual, and from the same allele in a homokaryotic individual, can form a heterodimer. For the experiment we chose the same alleles used in structural analyses. We synthesized and cloned *HD1* and *HD2* into pGAD-C1 (pCH312) and pGBD-C1 (pCH478) vectors, purchased from Genewiz (Germany)^61^ (Supplementary Table 4). The *Saccharomyces cerevisiae* strain PJ69-4a (CHY1268)^61^ (Supplementary Table 5) was co-transformed by lithium acetate transformation using every possible combination of *HD1* and *HD2*, with each *HD* alternately serving as either bait or prey in different runs of the experiment. We selected for transformants containing both plasmids by plating on SD -leu -trp and assessed the strength of protein-protein interactions using both the *HIS3* and *lacZ* reporter genes. *HIS3* expression was determined by plating on SD -leu -trp -his + 3AT and β-galactosidase activity of transformants was determined in triplicate using o-nitrophenyl-β-galactosidase as a substrate.

### Homokaryon identification using Sanger sequencing

To discover whether homokaryotic reproduction is common, we looked for homokaryotic sporocarps in populations of *A. phalloides* introduced to North America and from European populations. We aimed to identify heterozygotic SNP sites using Sanger sequencing. We used a collection of 109 sporocarps held in the Pringle laboratory fungarium, including 40 from invasive range (15 from three sites in and around Berkeley, California, 15 from a site in New Jersey, ten from a site in New York, eight from two sites in Canada) and 69 from native range (12 from a site Montpellier, France, six from two sites in Norway, 14 from 12 sites in the UK, 13 from four sites in Austria, two from two sites in Estonia, 12 from two sites in Hungary, and two from a site in Switzerland (Supplementary Table 6). In addition, we used a collection of 30 sporocarps collected from PRNS, California in 2021 for attempting to rediscover the homokaryotic individuals (Supplementary Table 2).

After extracting DNA from sporocarps, we PCR-amplified and Sanger sequenced beta-flanking genes using two pairs of newly designed primers (Supplementary Table 3 and Supplementary Methods). For samples without heterozygosity at the beta-flanking gene, we designed primers for and sequenced ten additional BUSCOs located on nine different contigs (Supplementary Table 3). Each target gene fragment had at least one SNP site with estimated allele frequency close to 0.5 in the population. We used the same amplification and sequencing methods as we used for the beta-flanking gene. Assuming the nine contigs are unlinked, and assuming each contig is itself strictly linked, we estimated the probability of misidentifying a heterozygotic sample as homozygotic to be lower than 0.002 (Supplementary Table 3).

### Statistics

The statistical tests used with yeast-two-hybrid data are described in the legend of Fig. 4.

## Data availability

Raw genomic reads used in this study are accessible in Sequence Read Archive (SRA) BioProject PRJNA565149. Raw transcriptomic reads have been deposited in SRA BioProject PRJNA689850.

## Code availability

Scripts and additional supporting information are available on Open Science Framework (https://osf.io/bq2ru/?view_only=9f17ee8694ff49858e0fa0b75a1854dc and https://osf.io/kde9c/?view_only=79b952e214ee4091a77067eed7e1a328).

