## Extended Data, will be used for the link to the file on the preprint site. for "Invasive Californian death caps develop mushrooms unisexually and bisexually"

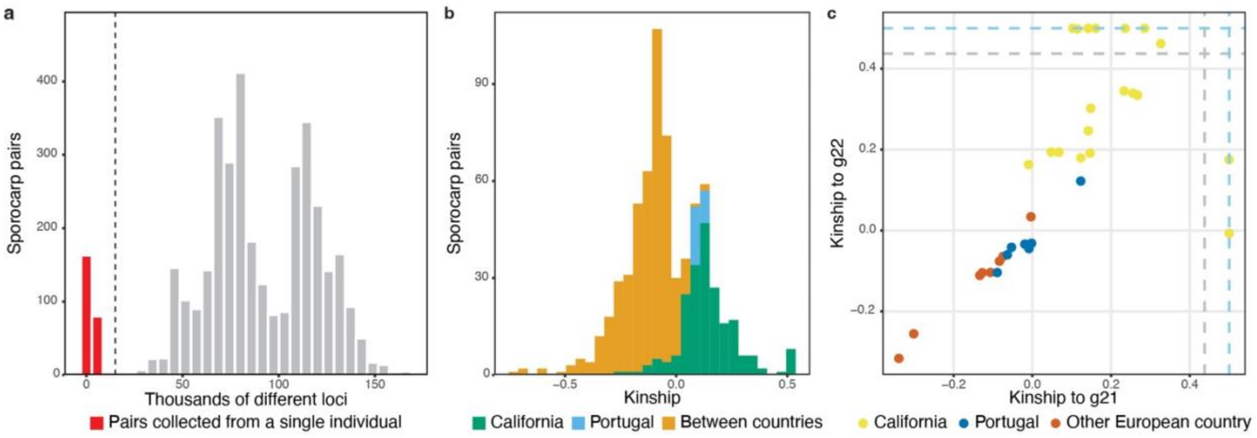


**Extended Data Fig. 1. Clone-correction and evidence for mating between homokaryotic individuals (g21 and g22) and heterokaryotic individuals.** (a) Numbers of SNPs differentiating pairs of sporocarps. Note distinct peak at left. Each of these sporocarp pairs represents two mushrooms collected from a single genetic individual. (b) Kinship estimates between pairs of individuals from California, pairs of individuals from Portugal, and between pairs of individuals collected in different countries (e.g. California-Portugal, Scotland-France, etc.). (c) Kinship estimates between heterokaryotic individuals and g21 or g22. Sky-blue dashed lines: kinship = 0.5. Grey dashed lines: kinship = 0.4375 (the threshold used to distinguish immediate kin (parents or offspring) from other kinds of relationships).

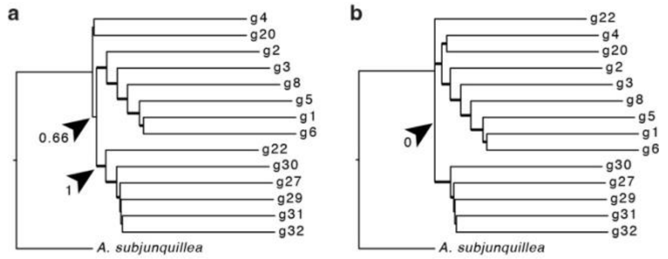


**Extended Data Fig. 2. Coalescent-based trees of *A. phalloides* individuals.** Phylogenies were reconstructed with ASTRAL unconstrained (a) or constrained by forcing g22 to be the outgroup of all other *A. phalloides* (b). Arrow heads mark branches leading to g22 (a) or the branch between g22 and other *A. phalloides* (b). Numbers indicate local posterior probabilities. There was no support for individual g22 as an outgroup for all other *A. phalloides* individuals, hence no support for the hypothesis of g22 (or g21) as a separate species. Branches with local posterior probabilities higher than 0.8 are thickened. Note the terminal branch lengths are forced to one in ASTRAL.

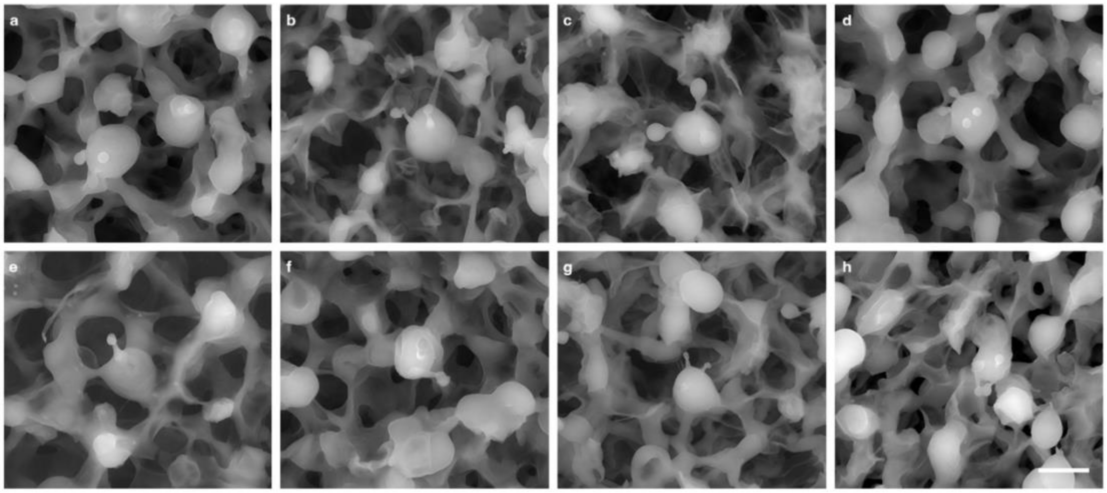


**Extended Data Fig. 3. Scanning electron microscopy of basidia.** (a–d) 1- to 4-spored basidia in heterokaryotic sporocarps. (e–h) 1- to 4-spored basidia in homokaryotic sporocarps. Scale bar: 10 µm.

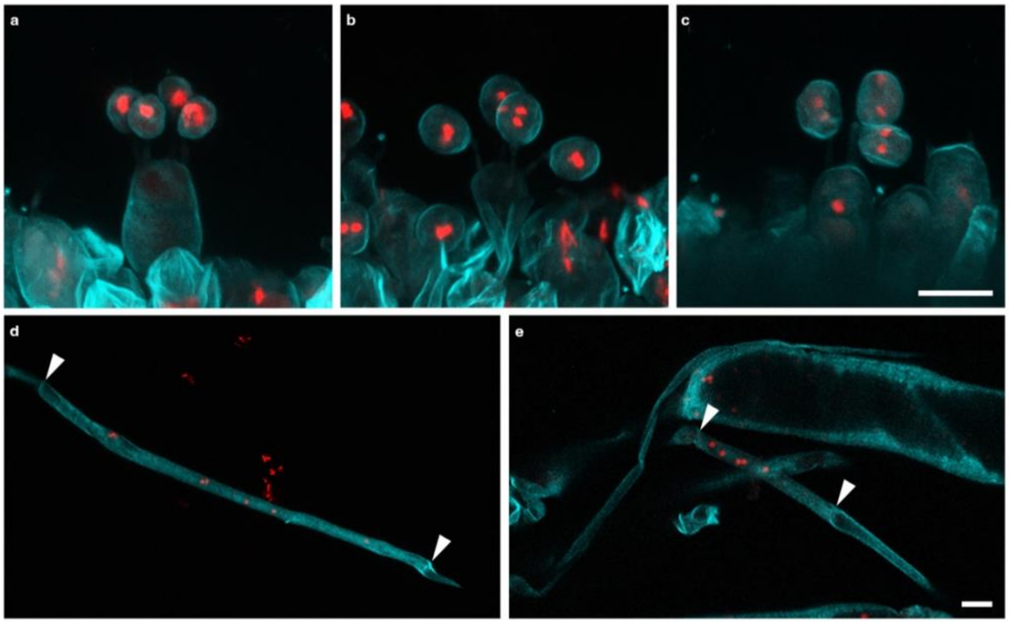


**Extended Data Fig. 4. Confocal microscopy of basidia and hyphae.** Composite images were created with Z-stack. (a–b) 4-spored basidia of heterokaryotic sporocarps. (c) 3-spored basidia of a heterokaryotic sporocarp. (d–e) Hyphae in the piths of stipes of heterokaryotic (d) and homokaryotic (e) sporocarps. Red: Vybrant Orange (nuclei); cyan: Calcofluor White (cell wall). Scale bar: 10 µm. Arrowheads: septae.

5

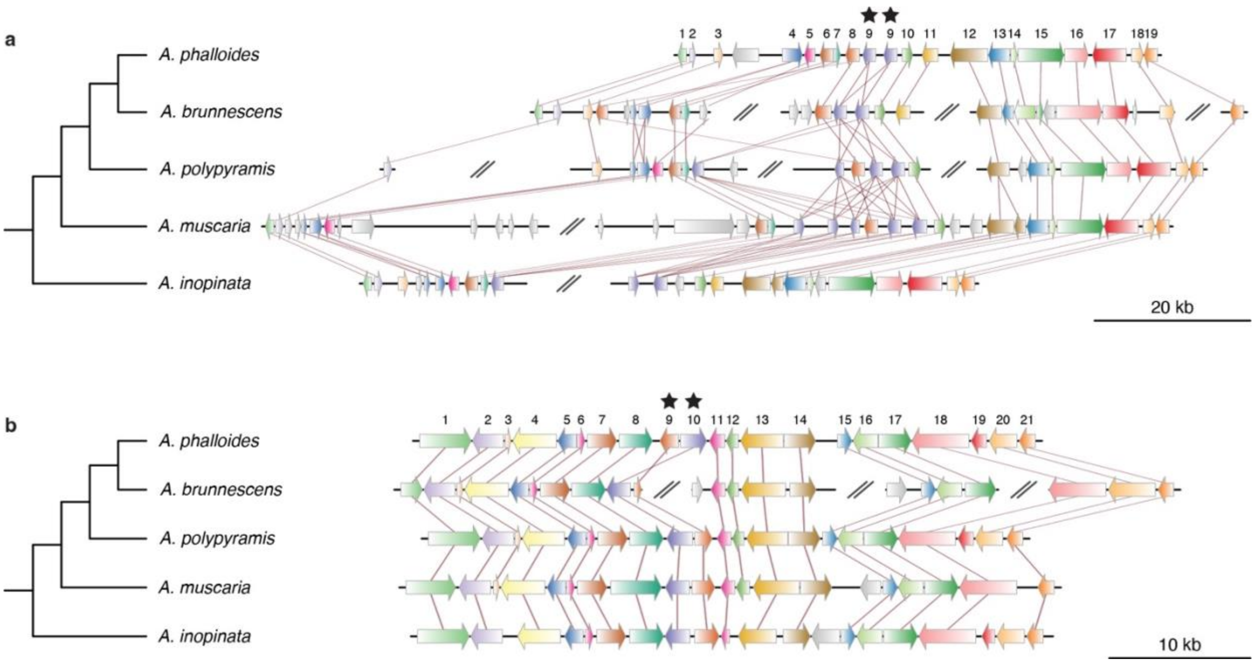
a
A. phalloides
A. brunnescens
A. polypyramis
A. muscaria
A. inopinata
1 2 3 4 5 6 7 8 9 9 10 11 12 13 14 15 16 17 18 19
20 kb
b
A. phalloides
A. brunnescens
A. polypyramis
A. muscaria
A. inopinata
1 2 3 4 5 6 7 8 9 10 11 12 13 14 15 16 17 18 19 20 21
10 kb

**Extended Data Fig. 5. Synteny of P/PR (a) and HD (b) loci across *Amanita* species.** The relatively strong synteny in the HD locus was consistent with findings from other species. Putative mating type determining genes (*PR*s and *HD*s) are labeled with stars. On the P/PR locus, twenty genes are homologous to genes in other *Amanita* species, and are functionally annotated as: (1) Cyanamide hydratase, (2) Hypothetical, (3) Hypothetical, (4) NUDIX hydrolase, (5) Nucleotide exchange factor Fes1, (6) Hypothetical, (7) Hypothetical, (8) Leucine-rich repeat domain superfamily, (9) GPCR fungal pheromone mating factor, (10) Hypothetical, (11) Cytochrome P450, (12) Proteasome non-ATPase regulatory subunit 13, (13) Hypothetical, (14) Prefoldin subunit 3, (15) Helicase superfamily 1/2, ATP-binding domain, (16) Meiotically up-regulated protein Msb1/Mug8, (17) UDP-Glycosyltransferase/glycogen phosphorylase, (18) Hypothetical, and (19) Eukaryotic translation initiation factor 2 subunit 3. On HD locus, twenty-one genes are homologous to genes in other *Amanita* species, and are functionally annotated as: (1) Glycine dehydrogenase, (2) Pentatricopeptidase repeat-containing protein, (3) Sec61p translocation complex subunit, (4) Hypothetical, (5) DUF1751, (6) DUF1754, (7) Hypothetical, (8) Mitochondrial intermediate peptidase, (9) HD2, (10) HD1, (11) Beta-flanking protein, (12) Hypothetical, (13) Glycosyltransferase family 8, (14) ABC1-domain-containing protein, (15) Mitotic spindle assembly checkpoint protein, (16) Nexin sorting protein, (17) Lung seven transmembrane receptor-domain-containing protein, (18) RPB2, (19) Fructosamine kinase, (20) MFS general substrate transporter, and (21) NADP dehydrogenase 1 alpha subcomplex subunit 9.

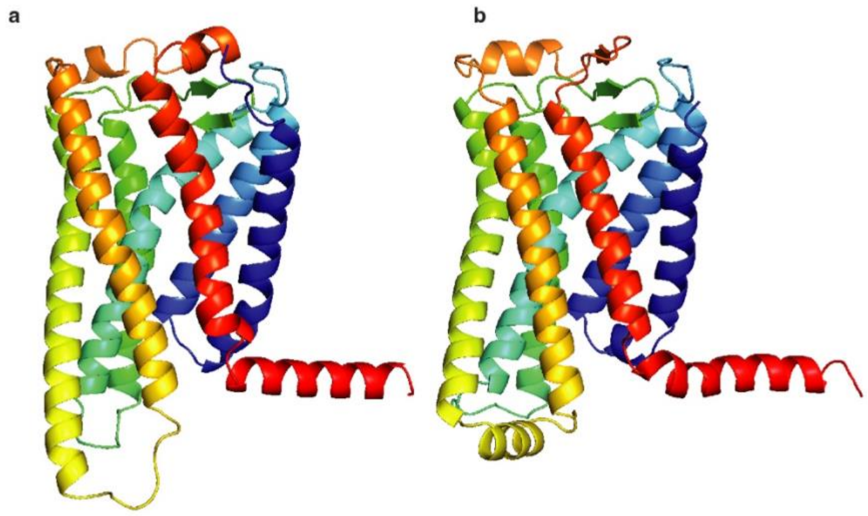


**Extended Data Fig. 6. Protein structures predicted for two putative *PR* genes.** Each shows similarity to the canonical structure of a GPCR (the protein family of *PR* genes): (a) *Ap.00g075660*, (b) *Ap*.00g075670. Low confidence regions are not shown.

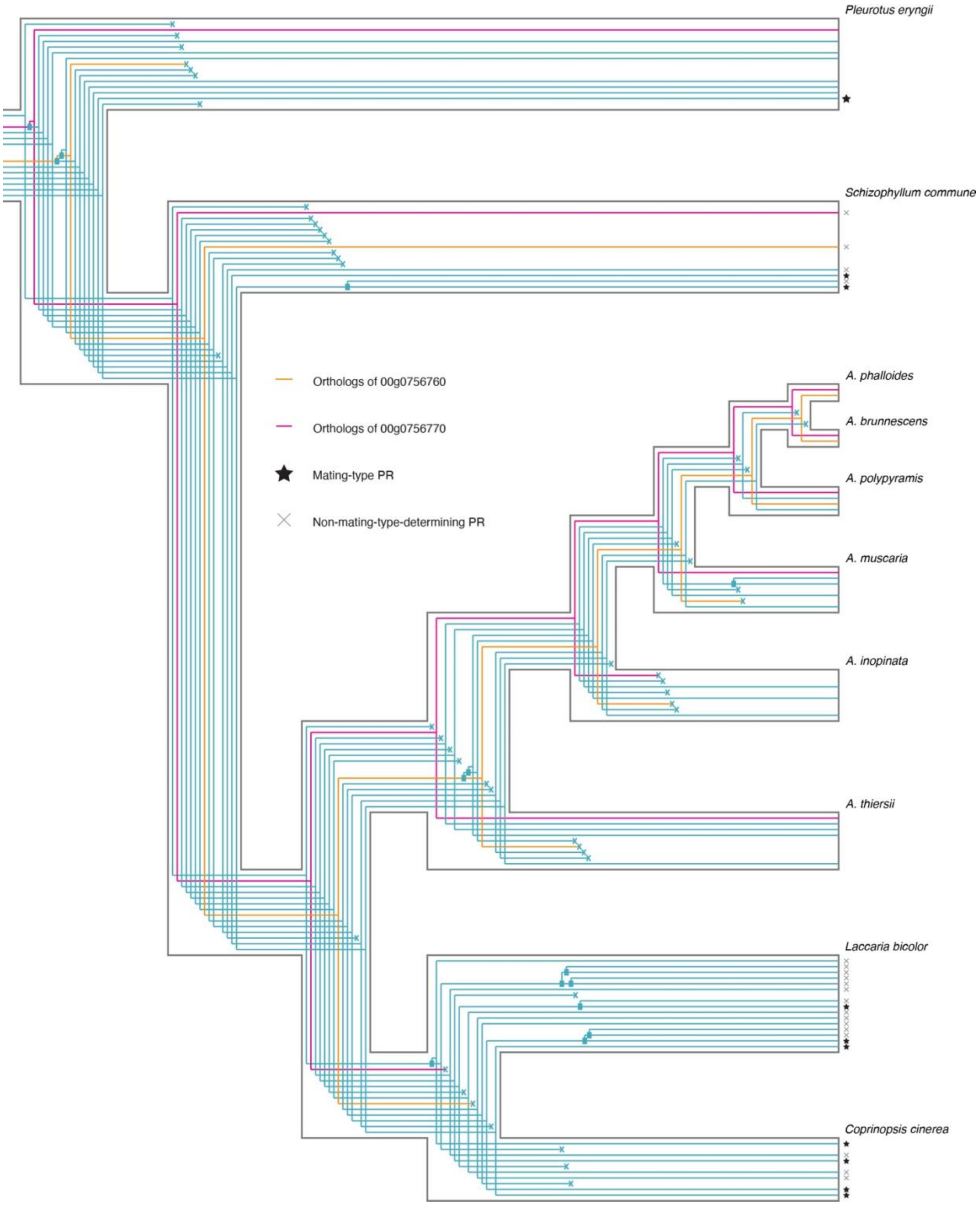
Pleurotus eryngii
Schizophyllum commune
A. phalloides
A. brunnescens
A. polypyramis
A. muscaria
A. inopinata
A. thiersii
Laccaria bicolor
Coprinopsis cinerea
Orthologs of 00g0756760
Orthologs of 00g0756770
Mating-type PR
Non-mating-type-determining PR

**Extended Data Fig. 7. Species-tree-aware protein phylogeny of PRs.** The phylogeny demonstrates orthology between PRs of *A. phalloides* and non-mating-type determining PRs from species of Agaricales. Note: older duplications may not indicate a duplication within a genome, instead, the duplication may represent a divergence between alleles in different mating types.

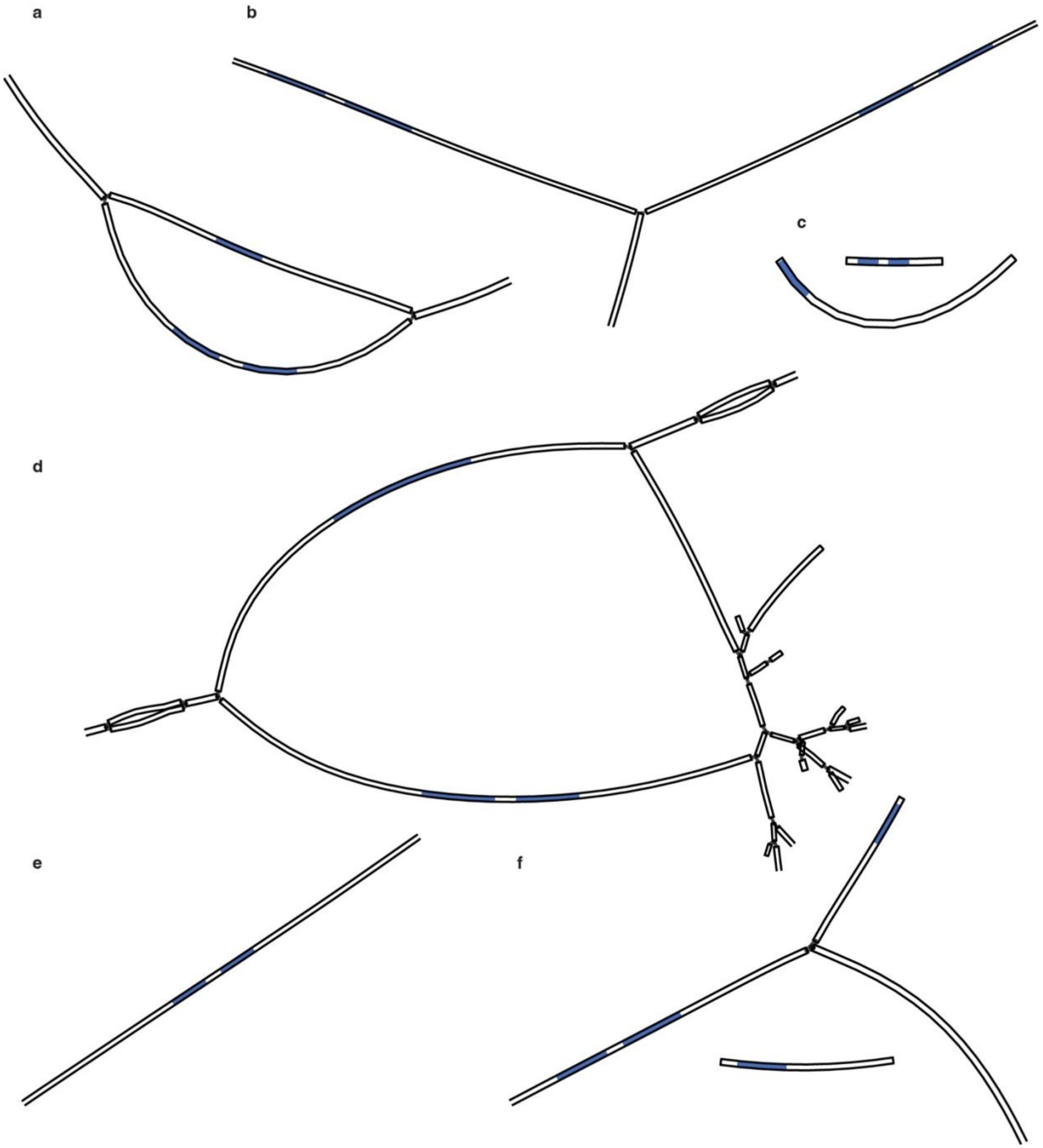


**Extended Data Fig. 8. Different assembly string graphs of HD locus.** (a) closed bubble (sample 10711), (b) open bubble (sample 10720), (c) detached (sample 10309), (d) complexed (sample 10511), (e) one unitig (sample 10303), and (f) three unitigs (sample 10240). Blue: BLAST hits with HD locus of reference genome as query.

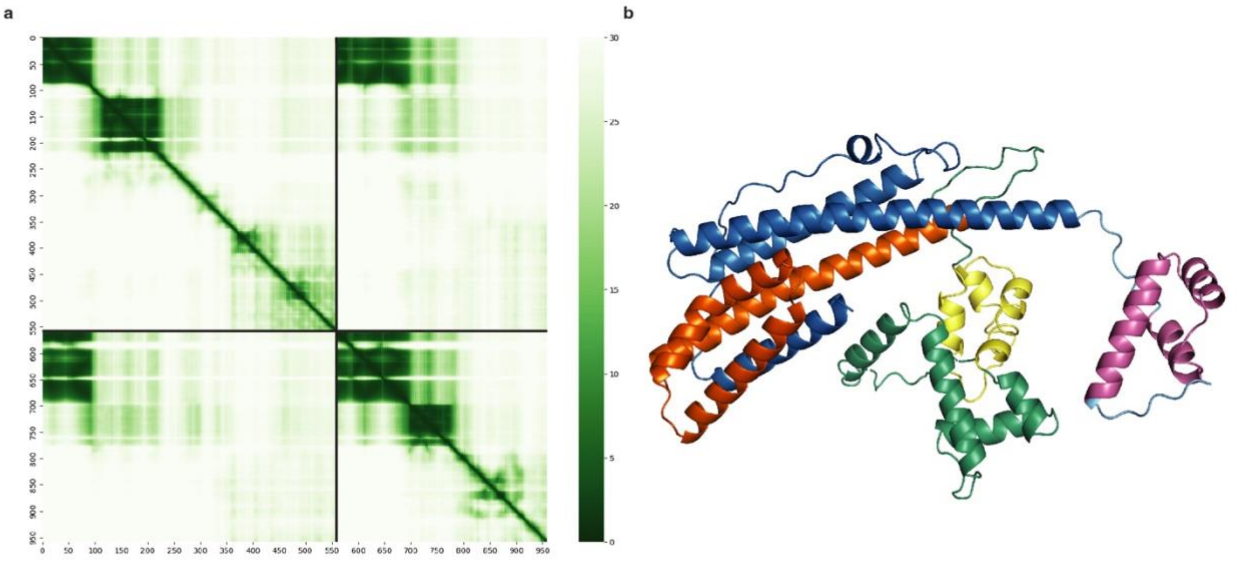


**Extended Data Fig. 9. Predicted heterodimer structures of HD1-8 and HD2-5.** N-terminals as binding sites and the two homeodomain motifs are highly supported. (a) Predicted alignment errors (PAE) between each pair of amino acids. HD1-8: 1–559; HD2-5: 560–959. Note the low PAE between N-terminals of the two proteins. (b) Cartoon of the heterodimer structure of HD1-8 (orange, green and yellow) and HD2-5 (blue, cyan and magenta). Low confidence regions are not shown. Orange and blue: putative binding domain; yellow and magenta: homeodomains; green and cyan: low confidence regions.

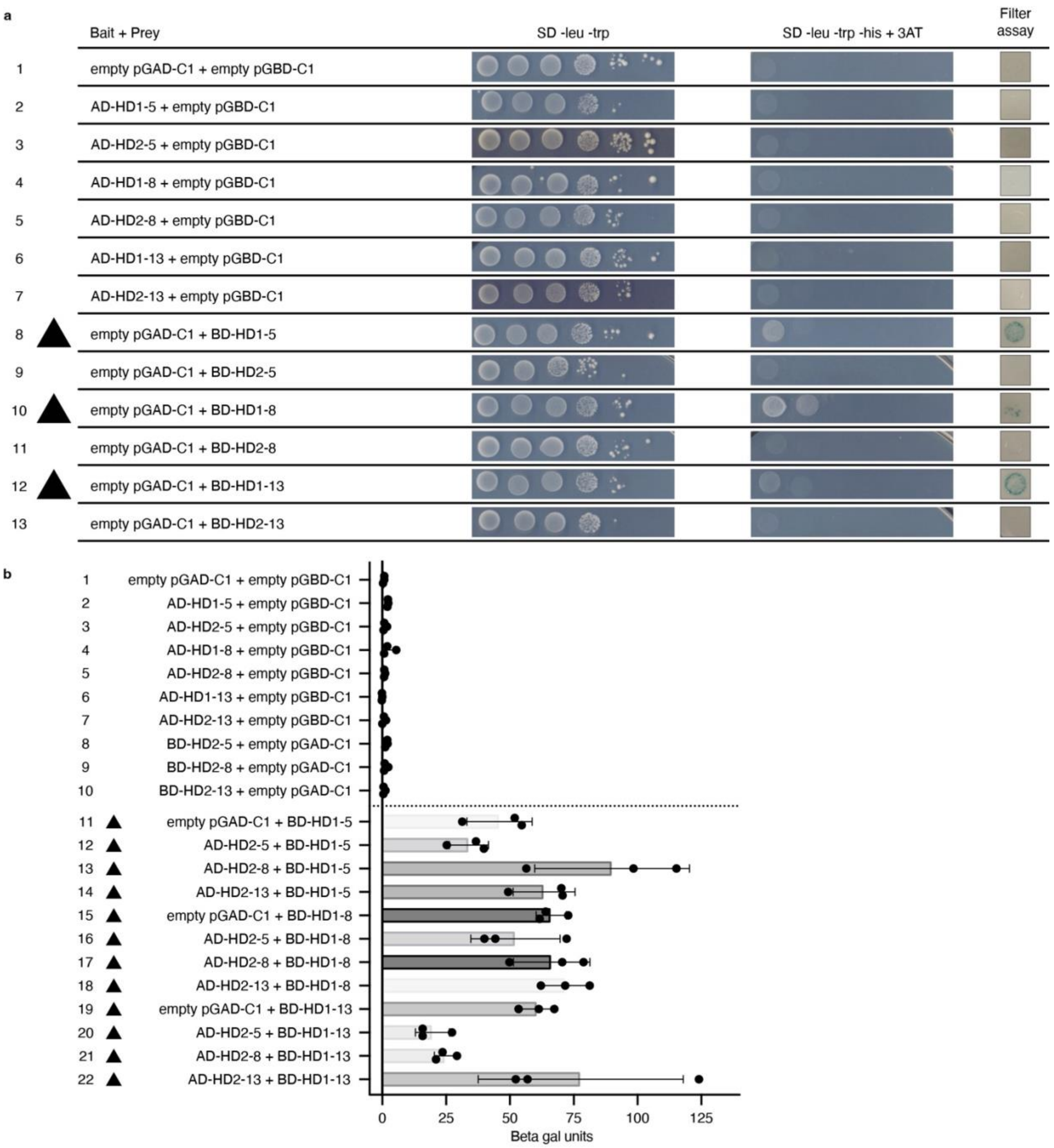
a
Bait + Prey
SD -leu -trp
SD -leu -trp -his + 3AT
Filter assay
1 empty pGAD-C1 + empty pGBD-C1
2 AD-HD1-5 + empty pGBD-C1
3 AD-HD2-5 + empty pGBD-C1
4 AD-HD1-8 + empty pGBD-C1
5 AD-HD2-8 + empty pGBD-C1
6 AD-HD1-13 + empty pGBD-C1
7 AD-HD2-13 + empty pGBD-C1
8 empty pGAD-C1 + BD-HD1-5
9 empty pGAD-C1 + BD-HD2-5
10 empty pGAD-C1 + BD-HD1-8
11 empty pGAD-C1 + BD-HD2-8
12 empty pGAD-C1 + BD-HD1-13
13 empty pGAD-C1 + BD-HD2-13
b
1 empty pGAD-C1 + empty pGBD-C1
2 AD-HD1-5 + empty pGBD-C1
3 AD-HD2-5 + empty pGBD-C1
4 AD-HD1-8 + empty pGBD-C1
5 AD-HD2-8 + empty pGBD-C1
6 AD-HD1-13 + empty pGBD-C1
7 AD-HD2-13 + empty pGBD-C1
8 BD-HD2-5 + empty pGAD-C1
9 BD-HD2-8 + empty pGAD-C1
10 BD-HD2-13 + empty pGAD-C1
11 empty pGAD-C1 + BD-HD1-5
12 AD-HD2-5 + BD-HD1-5
13 AD-HD2-8 + BD-HD1-5
14 AD-HD2-13 + BD-HD1-5
15 empty pGAD-C1 + BD-HD1-8
16 AD-HD2-5 + BD-HD1-8
17 AD-HD2-8 + BD-HD1-8
18 AD-HD2-13 + BD-HD1-8
19 empty pGAD-C1 + BD-HD1-13
20 AD-HD2-5 + BD-HD1-13
21 AD-HD2-8 + BD-HD1-13
22 AD-HD2-13 + BD-HD1-13
0
25
50
75
100
125
Beta gal units

**Extended Data Fig. 10. Yeast growth and reporter activity of control strains.** (a) Cells at the same starting concentration were 10-fold serially diluted and plated on selective plates. One representative transformant is shown for each interaction tested. Yeast growth was assessed on SD -leu -trp plates. Activity of the reporter gene *pGAL1-HIS3* was assessed by growth on SD -leu -trp -his + 3AT plates. Activity of the reporter gene *pGAL7-LacZ* was determined by filter assays and (b) liquid β-galactosidase assays. For each interaction, values for three independent biological replicates are shown (black dots). Each dot represents an average value of three technical replicates. Triangles represent reporter activity due to BD-HD1 auto-activation. Error bars represent standard error of the mean.
