## Supplementary Information, will be used for the link to the file on the preprint site. for "Invasive Californian death caps develop mushrooms unisexually and bisexually"

### Supplementary Methods

#### DNA extraction for genome sequencing

To extract DNA for genome sequencing, approximately 50 mg of cap tissue from each sporocarp was placed in a 2.0 ml microcentrifuge tube with four to five 3 mm glass beads and macerated using a MiniBeadbeater-8 (BioSpec Products Inc., Oklahoma) set at 75% speed for 1 min. 700 µl of CTAB buffer (2% cetyltrimethyl ammonium bromide, 2% PVP, 100 mM Tris-HCl, 20 mM EDTA, and 1.4 M NaCl [pH8.0]) was added after maceration, and samples were left to incubate at 60ºC for one hour. Next, 700 µl of a 24:24:1, by volume, phenol:chloroform:isoamyl alcohol solution was added to each sample and samples were gently mixed at room temperature for 10 min, followed by centrifugation at room temperature at 13,000 rpm for 10 min. The aqueous phase (~650–700 µl) of each sample was then carefully transferred to a new 2.0 ml tube. 700 µl of the phenol:chloroform:isoamyl alcohol solution was again added to each sample, and samples were inverted and mixed at room temperature for 10 min, followed by centrifugation at room temperature at 13,000 rpm for 10 min, at which point the aqueous phase was again transferred to a new 2.0 ml tube. Approximately 1.4 ml of 100% ethanol was added to each sample, and samples were incubated at -20ºC for 45 min, and then centrifuged at 4ºC at 13,000 rpm for 10 min. The supernatant was discarded, and the pellet dried in a Savant DNA 120 SpeedVac Concentrator (Thermo Fisher, Massachusetts) at room temperature for 10 min, or until dry, and finally resuspended in 400 µl of 10 mM Tris-HCl (pH 8.0) and transferred to a new 1.5 ml tube. To further purify genomic DNA, 12 µl of RNase A (Qiagen, Germany) was added to each sample, and each sample incubated at 37ºC for an hour. 16 µl of 5 M NaCl and 860 µl of 100% ethanol were then added to each tube and the solution left to precipitate at -20ºC for one hour, after which each tube was centrifuged at 4ºC at 13,000 rpm for 15 min and the supernatant was discarded. A final washing was performed with 500 µl of 75% ethanol; solutions were centrifuged at 4ºC at 13,000 rpm for 10 min and the supernatant discarded. Finally, the resulting pellet was resuspended in 200 µl of 10 mM Tris-HCl (pH 8.0). 5 ml of an Oxygen AxyPrep Mag PCR Clean-Up kit (Fisher Scientific, Pennsylvania) was used per manufacturer instructions to remove any remaining impurities. DNA was stored at -80ºC until it was provided to the University of Wisconsin-Madison Biotechnology Center.

**Genome library construction, sequencing and read filtering**

DNA was submitted to the University of Wisconsin-Madison Biotechnology Center. DNA concentration was verified using the Qubit® dsDNA HS Assay Kit (Life Technologies, Grand Island, NY). Samples were prepared according the TruSeq Nano DNA LT Library Prep Kit (Illumina Inc., San Diego, California, USA) with minor modifications. Samples were sheared using a Covaris M220 Ultrasonicator (Covaris Inc, Woburn, MA, USA), and were size selected for an average insert size of 550 bp using SPRI bead based size exclusion. Quality and quantity of the finished libraries were assessed using an Agilent DNA1000 chip and Qubit® dsDNA HS Assay Kit, respectively. Libraries were standardized to 2nM. Cluster generation was performed using the Illumina Rapid PE Cluster Kits v2 and the Illumina cBot. Paired-end, 251 bp sequencing was performed, using Rapid v2 SBS chemistry on an Illumina HiSeq2500 sequencer. Images were analyzed using the Illumina Pipeline, version 1.8.2.

Mean sequencing depth of each sample ranged from 10.56 to 150.86 (Supplementary Table 1; low depths characterized older specimens). Sequence data were filtered using Trim Galore! (ver. 0.4.5) (https://github.com/FelixKrueger/TrimGalore). Adapter trimming was set to the highest stringency so that even a single nucleotide of overlap with the adapter sequence was trimmed from a given read. After trimming, reads reduced to 100 bp or shorter and those with quality scores less than 30 were discarded.

To facilitate assembly of a high-quality reference genome, a sporocarp from Coimbra, Portugal (10511) was also sequenced on four SMRTcells on a PacBio RS II Sequel platform, also at the University of Wisconsin-Madison Biotechnology Center. PacBio sequencing resulted in raw coverage of 47x with an N50 read length of 6,310 bp.

**Reference genome assembly**

After testing five different genome assembly pipelines, we used an in-house hybrid approach to assemble the final reference genome. First, Illumina data of 10511 were subjected to a second round of filtering using Trimmomatic ver. 0.35[41] with the following parameters: ILLUMINACLIP:TruSeq3-PE-2.fa:2:30:10 CROP:245 LEADING:30 TRAILING:30 SLIDINGWINDOW:4:25 MINLEN:100. PacBio data were filtered to remove sequences shorter than 500 bp and error corrected with the Illumina data using FMLRC[62] and default settings. Error-corrected PacBio reads were then used to simulate 20x coverage 3 kbp insert size libraries with wgsim (https://github.com/lh3/wgsim) and parameter setting as follows: -e 0.0 -d 3000 -s 500 -1 100 -2 100 -r 0.0 -R 0.0 -S 123 -N 5000000. Illumina data and simulated long-range libraries were then assembled using AllpathsLG ver. 52400[63], setting HAPLOIDIFY=True. The resulting assemblies were subjected to further scaffolding using error-corrected PacBio data with the software LINKS v.1.8.5[64] with -d 2000,5000,10000,15000,20000 -t 20,20,5 and a k-mer value of 29. Scaffolds were extended and gap-filled using PBJelly v.15.8.24 and finally polished using Pilon v.1.2[65]. Polished assemblies were evaluated with QUAST[66] and BUSCO ver. 2 with the Basidiomycota database ver. 9[67]. The final assembly is 35.5 Mb, consisting of 605 scaffolds. The N50 and NG50 is 320 kbp and 184 kbp, respectively. The assembly encodes 1260 single-copy BUSCOs (94.4%).

**Variant calling and SNP filtering**

After the reference genome was assembled, single nucleotide polymorphisms (SNPs) and insertions and deletions (indels) in all genomes were identified using the Genome Analysis ToolKit (GATK) software v3.8-0-ge9d806836[29], following GATK best practices. Illumina reads from each of the 86 Illumina genome libraries were first mapped to the hybrid reference assembly using BWA-0.7.9[68] with the following parameters: mem -M -t 8 -v 2. Mapping rates for *A. phalloides* specimens ranged from 20.0% to 95.3%, with a median mapping percentage of 86.1% (only older specimens mapped at less than 50%). The mapping rate of 10511's Illumina reads to the 10511 hybrid assembly was 93.8%. Duplicate reads were marked, and the GATK program Haplotypecaller was used to call variants simultaneously on all samples, using MODE=DISCOVERY and type=GVCF. Due to the lack of known variants in *A. phalloides*, the raw variants were hard-filtered according to GATK's default parameter of the VariantFiltration program. The pipeline resulted in Variant Call Files (VCFs) containing 212,119 indels and 1,580,133 SNPs. To identify genetic individuals, we only used SNPs and refer to this VCF file as the "raw VCF".

To eliminate any SNPs called as the result of sequencing errors, the raw VCF was additionally filtered using VCFtools ver. 0.1.14[30]. All transposable elements, multi-allelic sites, and contigs with putative contamination were removed. First, transposable elements were identified using REPET ver. 2.5 and removed. Multi-allelic sites were both identified and removed, and putatively contaminated contigs were identified with CAT ver. 5.0.3 (https://github.com/dutilh/CAT; contigs 330, 313 and 581) and also removed. In addition, sites with a sequencing depth below a minimum depth of 85% of the per specimen mean depth, and above a maximum depth of $DP + 4\sqrt{DP}$ of the per specimen mean depth, were removed[69]. Next, variants were re-called based on allelic depth: if the ratio of allelic depths for a given site was between 0.25 and 0.75, that site was called as heterozygous (0/1), and ratios below and above were called homozygous reference (0/0) and homozygous alternate (1/1), respectively. We refer to the new VCF file as the “filtered VCF”.

**Transcriptome sequencing**

To sequence the transcriptome of *A. phalloides* for genome annotation, total RNA from sample 10721 was extracted using an RNeasy Mini Kit (Qiagen, Germany). We first macerated around 100 mg of tissue in 700 µl RLT Buffer and 7 µl β-mercaptoethanol with three 2.7 mm glass beads using a MiniBeadbeater-8 (BioSpec Products Inc., Oklahoma) for 1 min at 3000 rpm. We chilled the sample on ice for 1 min and macerated again for 1 min using the same settings. After centrifugation for 3 min at 14,000 rpm, 650 µl of supernatant was transferred to a new microcentrifuge tube. Then we added 650 µl of 100% ethanol and gently shook the tubes ten times to mix. We passed 650 µl of the solution through an RNeasy mini column with 14,000 rpm centrifugation for 15 sec twice. To clean up the RNA extract, the column was washed with 350 µL RW1 buffer with 14000 rpm centrifugation for 15 sec, and DNA was removed by incubating the column in 80 µL DNase solution (7 RDD buffer (Qiagen, Germany):1 DNase stock solution (Qiagen, Germany)) for 15 min, then washed with 350 µL RW1 buffer once and 500 µL RPE buffer twice with an additional 1.5 min centrifugation for the last wash. Finally, RNA was eluted with 30 µl RNase-free dd$H_2O$ by incubating for 2 min and 14,000 rpm centrifugation for 2 min twice. We collected the flow-through solution and stored it at -80°C prior to sequencing at the Biotechnology Center of the University of Wisconsin-Madison.

Purity and integrity of the total RNA was assessed via the NanoDrop One Spectrophotometer (ThermoFisher, Inc., Carlsbad, CA, USA) and Agilent 2100 Bioanalyzer (Agilent Technologies, Inc., Santa Clara, CA, USA), respectively. Stranded RNA libraries were prepared from samples that met the TruSeq™ Stranded Total RNA With Illumina® Ribo-Zero™ Plus rRNA Depletion input guidelines using the Illumina® TruSeq Stranded Total RNA with Ribo-Zero Plant kit (Illumina Inc., San Diego, California, USA). For each library preparation, cytoplasmic, mitochondrial and chloroplast ribosomal RNA was removed using biotinylated target-specific oligos combined with Ribo-Zero rRNA removal beads. Following purification, the RNA was fragmented using divalent cations under elevated temperature. Fragmented RNA was copied into first stranded cDNA using SuperScript II Reverse Transcriptase (Invitrogen, Carlsbad, California, USA) and random primers. Second strand cDNA was synthesized using a modified dNTP mix (dTTP replaced with dUTP), DNA Polymerase I, and RNase H. (The incorporation of dUTP quenches the second strand during amplification.) Double-stranded cDNA was cleaned up with AMPure XP Beads (1X) (Agencourt, Beckman Coulter). The cDNA products were incubated with Klenow DNA Polymerase to add a single ‘A’ nucleotide to the 3’ end of the blunt DNA fragments. Unique dual indexes (UDI) were ligated to the DNA fragments and cleaned up with two rounds of AMPure XP beads (0.8X). Adapter ligated DNA was amplified by PCR for 10 cycles and cleaned up with AMPure XP beads (0.8X). Final libraries were assessed for size and quantity using an Agilent DNA1000 Screentape and Qubit® 1X dsDNA HS Assay Kit (Invitrogen, Carlsbad, California, USA), respectively. Libraries were standardized to 2nM. Paired-end 2x150bp sequencing was performed, using standard SBS chemistry (v3) on an Illumina NovaSeq6000 sequencer. Images were analyzed using the standard Illumina Pipeline, version 1.8.2.

We received 170,345,047 paired-end raw reads of 126 bp sequences. The raw data were trimmed using Trimmomatic with tags “ILLUMINACLIP:TruSeq3-PE-2.fa:2:30:10”, “MAXINFO:30:0.4”, and “MINLEN:75” to remove adapters, short reads and unpaired reads. 142,428,127 read pairs were retained. As a preliminary assessment of data quality, we aligned the trimmed reads to the reference nuclear genome using HISAT2 ver. 2.1.0[70] setting a minimum intron length of 20 and a maximum intron length of 500. We observed an alignment rate of 81.5%.

**Reference genome annotation**

We annotated the reference genome with GenSAS server[28] using the transcriptome as evidence to train several gene predictors and using EvidenceModeler ver. 1.1.1[71] to subsequently weigh and combine the predictions. We first *de novo* assembled the transcriptome using Trinity ver. 2.2.0[72] with --jaccard-clip flag. In addition, we performed a second assembly of the transcriptome, guided by the genome, by mapping raw reads to the genome while limiting the maximal intron length at 300, using HISAT2 and assembling the mapped reads using Trinity with --jaccard-clip flag. Before undertaking the next steps of our annotation pipeline, we identified repetitive regions in the genome with RepeatMasker ver. 4.0.7[73] on GenSAS, using rmblast, quick speed and a fungal repeat database. We also used RepeatModeler ver. 1.0.11[74] on GenSAS to identify novel repetitive regions. The repetitive regions were then masked from the genome based on the outputs from both RepeatMasker and RepeatModeler. We then generated gene models from the combined *de novo* and genome-guided transcriptome assemblies with PASA[75] using default settings on GenSAS. We also generated a refined set of gene models by using the *exonerate* and *prepare_golden_genes_for_predictors.pl* tools from the JAMg[76] pipeline to pick the best gene models from the original set.

To *de novo* predict nuclear genes, we used five gene predictors: GeneMark-ES ver. 4.38[77], BRAKER ver. 2.1.0[78], SNAP[79], AUGUSTUS ver. 3.3.1[80] and CodingQuarry ver. 2.0[81]. We ran GeneMark-ES on GenSAS with default settings. We also ran BRAKER on GenSAS but trained it with the mapped transcriptome generated from HISAT2 on GenSAS. We trained SNAP and AUGUSTUS with the refined set of gene, and trained CodingQuarry with the original set.

To combine gene predictions, we tuned the weights of the five different predictors for EvidenceModeler based on: their individual coverage of the benchmarking universal single-copy orthologs (BUSCOs) of basidiomycetes (OrthoDB ver. 9)[67], gene boundaries, fragmentation/fusion and intron lengths of each prediction result and the different runs of EvidenceModeler. After exploring different weight combinations, we gave AUGUSTUS, BRAKER, GeneMark-ES, CodingQuarry, SNAP and PASA (transcript) weights of 5, 6, 6, 2, 5 and 10, respectively. Finally, we refined the gene models with PASA and used InterProScan ver. 5.29-68.0[82] and Pfam ver. 1.6[83] for functional annotation.

We identified 8,746 gene models with our pipeline, including 95.0% of basidiomycetes' BUSCOs (slightly different from the 94.4% resulting from our assembly of the reference genome). Our number of gene models was lower than a previous annotation from Pulman *et al.* (2016) (10,221 models), but our percentage of annotated BUSCOs was higher (in Pulman *et al.* (2016), 92% of BUSCOs were annotated)[84]. The difference between our annotation and Pulman *et al.*'s (2016) annotation may relate to assembly coverage. Our reference assembly was 35.5 Mb with an estimated 45.5 Mb genome (78% complete), whereas Pulman's assembly was 40 Mb (without an estimated genome size). Our annotation pipeline may also be more conservative than Pulman *et al*'s (2016) MAKER pipeline; when we used the BRAKER gene predictor (a combination of AUGUSTUS and MAKER) in our annotation pipeline, it identified 9,417 gene models. Finally, Pulman *et al.*'s (2016) genome was sequenced from California while our reference genome was collected in Portugal. The difference in gene model numbers may also reflect some degree of biological diversity. Nevertheless, estimates of BUSCO coverage suggest our genome annotation performs better on conserved genes, ideal for identification of conserved mating type loci.

#### Clone-correction

To understand which sequenced sporocarps were collected from a single genetic individual (from the same mycelium), we adapted methods from the R package *poppr*. Euclidean genetic distances were calculated between all pairs of sporocarps and visualized as a histogram (Extended Data Fig. 1a). A distinct peak in the histogram, near zero and apart from a second distinct peak, marks the sporocarps belonging to single individuals[85]. The 86 mushrooms resolve into 37 individuals, including 27 individuals consisting of only one mushroom and ten individuals consisting of multiple mushrooms (Extended Data Fig. 1a and Supplementary Table 1). No individuals spanned different collecting sites. We also used a second approach developed by us to identify kinship and confirmed the results generated using Euclidean genetic distances (see Methods).

#### Imaging

To compare the number of spores per basidium in homokaryotic and heterokaryotic *A. phalloides*, we imaged the basidia of five individuals (g21–g25, which are specimens of high quality) using a scanning electron microscope (SEM). We cut lyophilized gill tissue into pieces approximately 5 × 5 $mm^2$ in size and mounted tissue directly to a sample stub with conducting tape. We observed basidia under a FEI Quanta 200 SEM using the environmental SEM settings with a 25 kV electron beam with a spot size of 4.5 under 2.5 Torr, 5°C, 5000X magnification, and approximately 10 mm of working distance.

To understand the organization of nuclei within basidia, we first rehydrated and fixed lyophilized gill tissues in 10% neutrally buffered formalin for an hour at room temperature, followed by a single wash in dd$H_2O$. Then we stained tissues with 2 µg/ml Calcofluor White (Sigma-Aldrich, Missouri) in darkness for 20 min and subsequently washed the tissues with water. We hand-sliced gill thinly with a razor blade on a glass slide and stained them with a 5X Vybrant Orange dye (Thermo Fisher, Massachusetts) in darkness for 30 min under a coverslip.

We also sought to understand the organization of nuclei within homokaryotic and heterokaryotic mycelia, and so we cut the pith of fresh stipe tissues from samples collected in 2021 (20030 and 20053) and fixed the tissues as described above, later splitting the mycelia with forceps and scalpels. We stained the tissues with a freshly mixed dye containing 2 µg/mL Calcofluor White and 4X Vybrant Orange for 30 min in darkness under a coverslip.

To visualize nuclei of basidia and mycelia, slides were mounted under an LSM 710 confocal microscope (Zeiss, Germany) using a C-Apochromat 40X/1.20 W Korr objective lens. We used two channels to separate the fluorescent signals from Calcofluor White and Vybrant Orange. For Calcoflour White, we used a wavelength of 405 nm for excitation and detected the emission from 394 to 515 nm. For Vybrant Orange, we used a wavelength of 514 nm for excitation and detected the emission from 525 to 678 nm. The pinhole size was set to one airy unit for the longer wavelength. Because nuclei often cannot be captured on a single optical section, we performed z-stacking with a step size of 0.49 µm and constructed 2D images with maximum intensity projection using the Z project function in FIJI[86]. Slices were removed when bacterial contaminants overlap with hyphal compartments. Images were lastly cleaned up with denoise function in FIJI[86].

**DNA extraction and amplification for Sanger sequencing**

We extracted DNA by first grinding around 10 mg of tissue in 500 µl CTAB buffer (2% cetyltrimethyl ammonium bromide, 100 mM Tris-HCl, 20 mM EDTA, and 1.4 M NaCl [pH8.0]) and incubating samples at 65°C for 60 min. We gently mixed each solution with 500 µl 24:1 chloroform:isoamyl alcohol (Sigma-Aldrich, Missouri) for 5 min, centrifuged for 10 min at 12,000 rpm in a 5417C centrifuge (Eppendorf, Germany), and moved supernatants to new tubes, repeating these steps twice. We then incubated supernatants with 0.6X volume of isopropanol in a -20 ºC freezer overnight. We centrifuged each solution for 7 min at 12000 rpm and removed the supernatant. The DNA pellets were washed with 1 ml ice cold 70% ethanol and centrifuged for 2 min at 12,000 rpm twice. Ethanol was discarded and the DNA pellets were dried in a Savant DNA 120 SpeedVac Concentrator (Thermo Fisher, Massachusetts) for 30 min. Finally, we eluted DNA with 50 µl water and stored the DNA extract at -20°C.

We amplified fragments of the variable beta-flanking gene of the HD locus for the first heterozygosity screen using a C1000 Touch Thermo Cycler (Bio-Rad, California). We originally designed a primer pair to sequence a 691-mer region, but because we failed to amplify this region from many of our older specimens, we later designed a primer pair with an amplicon size of 248 nucleotides (Supplementary Table 3). For each PCR reaction, a final volume of 25 µl of reagents contained 1 µl of DNA template, 1X EconoTaq PLUS Master Mix (Lucigen, Wisconsin), 0.4 µM forward and 0.4 µM reverse primers. The PCR cycle included an initial denaturation at 95°C for 7 min, 35 rounds of denaturing at 95°C for 15 sec, annealing at 55°C for 30 sec and elongation at 72°C for 1 min, and additional 7 min of elongation at the end of the cycles. The PCR products were sequenced by Functional Biosciences (Wisconsin) on ABI 3730xl instruments (Thermo Fisher, Massachusetts).

#### Supplementary Discussion

##### Explanations for the absence of heterozygosity

The simplest explanation for an absence of heterozygosity is homokaryosis: individuals house the same haploid genome in every nucleus of the mycelium (either as a single copy (n) or as two copies (2n)). If we are wrong, and some nuclei house one copy of the genome while others house two (i.e. a mix of haploid and diploid nuclei), then the haploid genome would be considered as the unit driving observed phenomena, and statements like "nuclei appeared to have persisted in invaded habitats for at least 17 years" would translate to "haploid genomes appeared to have persisted in invaded habitats for at least 17 years".

##### Other strategies to identify mating systems

Traditionally, mating systems of fungi are explored with mating experiments, by co-culturing sexual spores from sporocarps and observing subsequent behaviors. However, we cannot adopt this strategy because *A. phalloides* cannot be cultured.

##### The natural history of the homokaryon g22 does not support an hypothesis of pseudosexuality

Fungal sexual systems are diverse and intensively studied. We did consider a range of alternative hypotheses to explain our data, but none appear to describe the dynamics we observe both in nature and the laboratory. In particular, we considered the hypothesis of pseudosexuality[22]. But pseudosexuality is rare (to date described from a single species), and in the species where it is described it is also rare (only 1% of basidia). The rarity would suggest that if pseudosexuality is found in *A. phalloides*, individual g22 is the result of a single pseudosexual event. Because we find this individual at both our Drake 2 and Drake 4 sites, and because *A.*

*phalloides* does not produce asexual spores[87], pseudosexuality as an explanation would require an enormous mycelium more than 200 m in diameter. This size is inconsistent with our data on the sizes of *A. phalloides* individuals in both Europe and North America (Pringle unpublished). Pseudosexuality does not emerge as a likely explanation for the discovery of homokaryotic *A. phalloides* sporocarps.

#### Supplementary Notes

##### Heterozygosity estimation, k-mer analysis and allele frequency

The estimated heterozygosities of most individuals are normally distributed with a range between $1.7 \times 10^{-3}$ and $3.7 \times 10^{-3}$, but two outliers have estimated heterozygosities of a different order of magnitude, at $3.07 \times 10^{-4}$ (g21) and $2.5 \times 10^{-4}$ (g22). Both of these individuals are from Drake2, PRNS, California. Two sporocarps of individual g21 were found in 2014. One sporocarp of individual g22 was found in 2004 and five were found in 2014. (We collected both g21 and g22 again in 2021, but none of the genomes of the 2021 specimens were sequenced.)

Most heterozygotic individuals with a sequencing depth higher than 50x exhibit a typical pattern of k-mer and allele frequency distributions: in a k-mer frequency distribution, heterozygous individuals have a minor peak at half of the depth of major peak. In an allele frequency distribution, heterozygous individuals have a peak at 0.5. The most deeply sequenced sporocarps of the two individuals with low estimated heterozygosities have sequencing depths of 71.1x (g21) and 67.6x (g22), but neither have typical k-mer or allele frequency distributions, additional evidence they are homokaryotic. While four other individuals (g9, g10, g11 and g12) with sequencing depths higher than 50x also do not have visible minor peak in k-mer frequency distributions, the absence of minor peak in these individuals is likely caused by greater sequencing error: the frequencies of k-mers at 0.5 normalized k-mer depth of these four individuals were higher than frequencies of g21 and g22 (Fig. 1b).

In allele sequencing frequency plots, all heterozygotic individuals, including the four individuals without visible minor peaks in k-mer graphs, show a peak at 0.5. By contrast, the two individuals with low estimated heterozygosities display no peak, again suggesting they are homokaryotic (Fig. 1c). The allele sequencing frequency plots using the filtered VCF without the re-calling of variants generated nearly identical results to the plots generated using the filtered VCF with re-calling.

**Kinship analysis**

Two genetic individuals (four mushrooms collected in 2004 and 2015) are the parent or offsprings of g21 and six individuals (made up of 22 mushrooms collected in 1993, 2004, 2014, 2015) are the parent or offsprings of g22 (Extended Data Fig. 1b,c). Because only one individual can be the parent of a homokaryotic individual, both g21 and g22 must be mating. We assume the population is not intensively inbreeding: significant inbreeding would cause a significant reduction in heterozygosity, which was absent in the individuals identified as parent or offsprings (Fig. 1 and Supplementary Table 1). We note that while heterokaryotic individual g24 and the homozygous g22 share a high kinship (0.462), because most other heterokaryotic parent/offsprings of homokaryotic individuals had much higher estimated kinships, we took a conservative approach and do not consider g24 as a parent/offspring of g22.

**Pheromone identification**

Two hypothetical genes encoding putative pheromones were identified on two contigs (4 and 135) and both had conserved -CaaX motifs (Supplementary Fig. 1). However, only one hypothetical pheromone precursor had the ER/DR motif commonly found in pheromone precursors of model species[21]. The phenomenon is also observed in close relatives of the death cap, including *A. muscaria* var. *guessowii* and *V. volvacea*, which also do not have any ER/DR motifs in any putative pheromone precursors[88,89].

***De novo* genome assemblies used to identify HD genes at intraspecies level**

In the genome assemblies of most sporocarps there were two HD unitigs, but two and eleven sporocarps had three and one HD unitigs, respectively (Supplementary Table 1). The genomes of the two sporocarps with three HD unitigs have a low sequencing coverage (34.5x and 9.5x). Both specimens belong to heterokaryotic, multi-sporocarp genetic individuals, and other sporocarps of the same individuals possess two HD unitigs. The eleven samples with one HD unitig included all eight homokaryotic sporocarps, and three other putatively heterokaryotic sporocarps (10019, 10169 and 10233), each with a very low coverage (12.0–26.2x). HD unitigs of 10019 and 10169 were very short: 1,892 and 495 bp, whereas the HD unitig of 10233 was longer: 24,659 bp. Further mapping of raw reads back to the HD assemblies of 10019 and 10233 show heterozygosity at the locus, but we could not map raw reads back to the 10169 assembly. But we did not detect any heterozygosity at any HD locus in any homokaryotic sporocarp. We consider the results for 10019, 10169 and 10233 as caused by shallow sequencing.

Remaining sporocarps present two HD unitigs: 35, 28 and five of their genome assemblies form closed, open and complexed bubbles, respectively, and the two HD unitigs of the other five samples were detached (Supplementary Table 1). Specimens belonging to the same genetic individual can present differently, for example, three genomes with detached HD unitigs were assembled into closed, open or complexed bubbles in other sporocarps of the same genetic individuals. Two HD unitigs representing different alleles of the HD locus appeared to be present in most if not all individuals.

**Annotation of HD genes**

Annotations of the *HD* genes of sporocarps collected between 2004 and 2015 from California and Portugal identify 22 unique *HD1* alleles and 21 unique *HD2* alleles, which translate to 20 and 18 unique protein products, respectively. We excluded specimens 10233 and

10288 from analyses because of problems with the genome assemblies for the two specimens. The lengths of HD1 and HD2 proteins range from 532 to 563, and from 385 to 406, amino acids. The genes contain two and three introns, respectively, with intron lengths of 48 to 62, and 45 to 69, bp.

**Protein structural analysis**

PR proteins are membrane proteins, which usually have signal peptides, and belong to the GPCR family which is characterized by seven transmembrane helices. Both of *A. phalloides*' PR proteins encoded seven transmembrane helices, but neither of them had signal peptides. Instead of signal peptides, they may encode signal anchor sequences for translocating the proteins to membrane[90]. Structural models from AlphaFold2 also support a folding of the transmembrane domains into a tertiary structure resembling the structure of a typical GPCR (Extended Data Fig. 6).

In the HD1 from each allele, we identified a NLS at 310–360 amino acid, but no NLS was predicted in any HD2. Homeodomains were found at 100–175 amino acid in HD1 and 140–200 amino acid in HD2. The tertiary homeodomain structures predicted by AlphaFold-Multimer exhibited a high structural similarity with the mating type HD of *Saccharomyces cerevisiae*. AlphaFold-Multimer also predicted low predict alignment errors (PAEs) among amino acids in the N-terminal of the two proteins suggesting a high confidence of them being binding sites for heterodimerization (Extended Data Fig. 9). However, the two homeodomains do not fit the quaternary structure of the mating type HD of *Saccharomyces cerevisiae* (UniProt Acc.: 1le8), probably because of the low confidence/flexible linkers between the N-terminal domains and homeodomains.

**Interspecies phylogeny of *PR* genes**

The species-tree-aware gene phylogeny suggests the two PRs in *A. phalloides* are orthologous to non-mating-type PRs in other agaricomycetes (Extended Data Fig. 7). However, the orthologs in non-agaricomycete species appear to be mating-type determining pheromone receptors. The different functions of these genes in different classes of fungi may be the result of a duplication of the two pheromone receptors at the ancestral branch of agaricomycetes, followed by neofunctionalization in the new gene copies.

**Yeast two hybrid of *HD* genes**

The ORFs of the *HD1* and *HD2* alleles 5, 8, and 13 were fused to the *GAL4* activation domain (AD) or *GAL4* DNA binding domain (BD) creating either “prey” or “bait” constructs, which were transformed into *S. cerevisiae* in all possible combinations. Interactions were determined by assessing multiple reporter genes, both qualitatively using filter assays and selective media (Extended Data Fig. 10a), and quantitatively using liquid β-galactosidase assays (Extended Data Fig. 10b). We found that HD1 from each allele when fused to the Gal4 BD resulted in autoactivation and expression of the reporters (Extended Data Fig. 10a, lines 8, 10, 12, and S11B, lines 11, 15, 19). Furthermore, addition of an HD2 fused to the Gal4 AD did not increase reporter expression levels in these strains (Extended Data Fig. 10b, lines 11-22), suggesting that potential interactions between the bait and prey proteins could be obscured by the BD-HD1 background activation. In contrast, HD1s fused to the Gal4 AD did not autoactivate the reporters (Extended Data Fig. 10a, lines 2, 4, 6) and were therefore used to assess interactions with HD2s fused to the Gal4 BD.

To test the possibility that the HD1 or HD2 from allele 13 could auto-interact and form homodimers to regulate homokaryotic sexual development, we tested all combinations of BD-

HD1 + AD-HD1 alleles and BD-HD2 + AD-HD2 alleles (Supplementary Fig. 2). As before, all BD-HD1 alleles were auto-activating and did not appear to change in the presence of any AD-HD1 alleles (Supplementary Fig. 2, lines 1-9), precluding an assessment of possible HD1-HD1 interactions. In contrast, we found that no combinations of BD-HD2 + AD-HD2 alleles induced expression of the reporter genes, indicating that HD2 of alleles 5, 8, and 13 are unlikely to form regulatory homodimers.

### Supplementary Figures

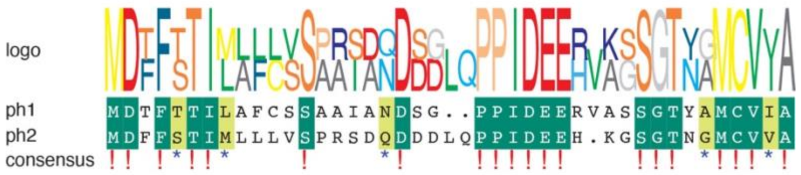

**Supplementary Fig. 1. Putative pheromone precursors.** The two pheromones both show the presence of the -CxxA motifs, characteristic of fungal pheromones. Neither gene was found near a *PR* gene.

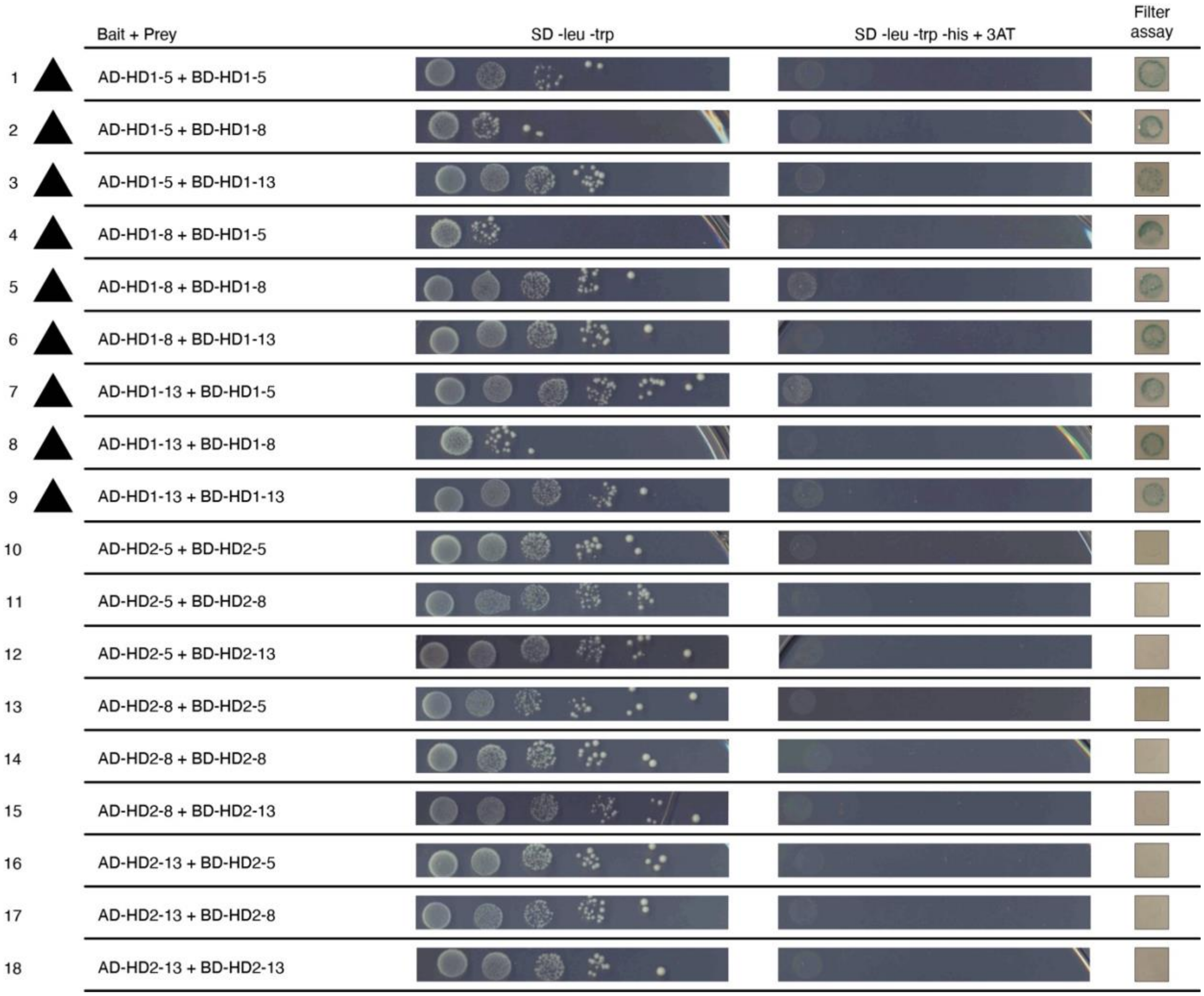

**Supplementary Fig. 2. Yeast growth and reporter activity of potential homodimer interactions.** Cells at the same starting concentration were 10-fold serially diluted and plated on selective plates. One representative transformant is shown for each potential interaction. Yeast growth was assessed on SD -leu -trp plates. Activity of the reporter gene *pGAL1-HIS3* was assessed by growth on SD -leu -trp -his + 3AT plates. Activity of the reporter gene *pGAL7-LacZ* was determined by filter assays. Triangles represent reporter activity due to BD-HD1 auto-activation.

### Supplementary Tables

#### Supplementary Table 1. Specimens with whole genome sequencing data

| ID | Genotype | Year | Latitude | Longitude | Country | Region | Collection site | Sequencing depth | Heterozygosity (‰) | HD assembly | HD1 allele* | HD2 allele* |
|---|---|---|---|---|---|---|---|---|---|---|---|---|
| 10004 | g1 | 1978 | 59.8586 | 17.6389 | Sweden | Uppland | NA | 57.4139 | 2.5543 | detached | NA | NA |
| 10007 | g2 | 1991 | 56.7044 | -3.7297 | Scotland | East Perthshire | NA | 60.6030 | 2.8408 | closed bubble | NA | NA |
| 10016 | g3 | 2000 | 52.3614 | 0.4866 | England | West Sussex | NA | 89.2649 | 3.6402 | closed bubble | NA | NA |
| 10018 | g4 | 2006 | 40.9211 | 9.1930 | Italy | Sardinia | NA | 94.6680 | 3.4372 | closed bubble | NA | NA |
| 10019 | g5 | 2000 | 54.4795 | -7.7315 | Northern Ireland | Fermanagh | NA | 26.2160 | 2.9201 | 1 | NA | NA |
| 10169 | g6 | 2003 | 55.9167 | 9.5333 | Denmark | Zealand | NA | 12.0349 | 2.4864 | 1 | NA | NA |
| 10170 | g7 | 1993 | 38.0525 | -122.8528 | USA | CA | NA | 45.4890 | 2.1329 | detached | NA | NA |
| 10171 | g8 | 2006 | 49.8109 | 14.9282 | Czech Republic | Central Bohemia | NA | 96.2488 | 3.7932 | closed bubble | NA | NA |
| 10221 | g9 | 2004 | 38.0548 | -122.8332 | USA | CA | Drake 2 | 53.0130 | 2.5699 | closed bubble | 13/15.2 | 13/15 |
| 10222 | g10 | 2004 | 38.0548 | -122.8332 | USA | CA | Drake 2 | 91.4194 | 2.8697 | closed bubble | 13/16.2 | 13/16.2 |
| 10223 | g11 | 2004 | 38.0548 | -122.8332 | USA | CA | Drake 2 | 71.9090 | 2.7636 | closed bubble | 13/15.1 | 13/15.1 |
| 10224 | g22 | 2004 | 38.0548 | -122.8332 | USA | CA | Drake 2 | 71.0926 | **0.2423** | 1 | 13 | 13 |
| 10225 | g12 | 2004 | 38.0548 | -122.8332 | USA | CA | Drake 2 | 56.8688 | 2.6853 | closed bubble | 1/13 | 1/13 |
| 10226 | g23 | 2004 | 38.0548 | -122.8332 | USA | CA | Drake 2 | 51.5766 | 2.6498 | open bubble | 8/13 | 8/13 |
| 10227 | g13 | 2004 | 38.0548 | -122.8332 | USA | CA | Drake 2 | 37.8999 | 1.6869 | open bubble | 11/- | 11/- |
| 10228 | g13 | 2004 | 38.0548 | -122.8332 | USA | CA | Drake 2 | 20.8024 | 1.6621 | detached | 11/- | 11/- |
| 10229 | g14 | 2004 | 38.0548 | -122.8332 | USA | CA | Drake 2 | 29.1773 | 2.5015 | closed bubble | 10/13 | 10/13 |
| 10230 | g15 | 2004 | 38.0548 | -122.8332 | USA | CA | Drake 2 | 58.7555 | 2.2826 | open bubble | 8/11 | 8/11 |
| 10231 | g16 | 2004 | 38.0548 | -122.8332 | USA | CA | Drake 2 | 22.8035 | 2.6475 | closed bubble | 11/13 | 11/13 |
| 10232 | g17 | 2004 | 38.0548 | -122.8332 | USA | CA | Drake 2 | 33.2382 | 2.4454 | open bubble | 10/13 | 10/13 |
| 10233 | g18 | 2004 | 38.0548 | -122.8332 | USA | CA | Drake 2 | 17.9585 | 2.1513 | 1 | -/- | -/- |
| 10237 | g19 | 2004 | 38.0552 | -122.8341 | USA | CA | Drake 3 | 41.9854 | 2.1915 | complexed | 8/13 | 8/13 |
| 10238 | g19 | 2004 | 38.0552 | -122.8341 | USA | CA | Drake 3 | 67.3885 | 2.3481 | detached | 8/13 | 8/13 |
| 10239 | g19 | 2004 | 38.0552 | -122.8341 | USA | CA | Drake 3 | 92.8748 | 2.8940 | complexed | 8/13 | 8/13 |

| | | | | | | | | | | | | |
|---|---|---|---|---|---|---|---|---|---|---|---|---|
| 10240 | g19 | 2004 | 38.0552 | -122.8341 | USA | CA | Drake 3 | 34.4694 | 2.2578 | 3 | 8/13 | 8/13 |
| 10241 | g19 | 2004 | 38.0552 | -122.8341 | USA | CA | Drake 3 | 58.0814 | 2.2726 | open bubble | 8/13 | 8/13 |
| 10277 | g20 | 2005 | 42.0396 | 9.0129 | France | Corsica | NA | 99.4189 | 3.4721 | closed bubble | NA | NA |
| 10280 | g23 | 2014 | 38.0548 | -122.8332 | USA | CA | Drake 2 | 48.3414 | 2.5836 | open bubble | 8/13 | 8/13 |
| 10281 | g23 | 2014 | 38.0548 | -122.8332 | USA | CA | Drake 2 | 85.3952 | 3.1884 | open bubble | 8/13 | 8/13 |
| 10282 | g23 | 2014 | 38.0548 | -122.8332 | USA | CA | Drake 2 | 45.2683 | 2.5994 | open bubble | 8/13 | 8/13 |
| 10283 | g23 | 2014 | 38.0548 | -122.8332 | USA | CA | Drake 2 | 56.9446 | 2.9749 | open bubble | 8/13 | 8/13 |
| 10287 | g24 | 2014 | 38.0548 | -122.8332 | USA | CA | Drake 2 | 33.0468 | 2.5622 | closed bubble | 13/15.1 | 13/15 |
| 10288 | g24 | 2014 | 38.0548 | -122.8332 | USA | CA | Drake 2 | 50.4221 | 2.6489 | closed bubble | 13/15.1 | 13/15 |
| 10292 | g24 | 2014 | 38.0548 | -122.8332 | USA | CA | Drake 2 | 76.7689 | 3.0443 | closed bubble | 13/15.1 | 13/15 |
| 10293 | g21 | 2014 | 38.0548 | -122.8332 | USA | CA | Drake 2 | 24.9338 | **0.4236** | 1 | 15.1 | 15.1 |
| 10294 | g24 | 2014 | 38.0548 | -122.8332 | USA | CA | Drake 2 | 79.7589 | 3.0396 | closed bubble | 13/15.1 | 13/15 |
| 10295 | g22 | 2014 | 38.0548 | -122.8332 | USA | CA | Drake 2 | 48.7380 | **0.2442** | 1 | 13 | 13 |
| 10298 | g21 | 2014 | 38.0548 | -122.8332 | USA | CA | Drake 2 | 67.6523 | **0.1730** | 1 | 15.1 | 15.1 |
| 10299 | g22 | 2014 | 38.0548 | -122.8332 | USA | CA | Drake 2 | 67.3467 | **0.2277** | 1 | 13 | 13 |
| 10300 | g22 | 2014 | 38.0548 | -122.8332 | USA | CA | Drake 2 | 61.0121 | **0.2229** | 1 | 13 | 13 |
| 10301 | g22 | 2014 | 38.0548 | -122.8332 | USA | CA | Drake 2 | 29.1512 | **0.3549** | 1 | 13 | 13 |
| 10303 | g22 | 2014 | 38.0548 | -122.8332 | USA | CA | Drake 2 | 56.4730 | **0.2005** | 1 | 13 | 13 |
| 10304 | g23 | 2014 | 38.0548 | -122.8332 | USA | CA | Drake 2 | 64.4879 | 3.1543 | open bubble | 8/13 | 8/13 |
| 10306 | g23 | 2014 | 38.0548 | -122.8332 | USA | CA | Drake 2 | 30.9192 | 2.6376 | open bubble | 8/13 | 8/13 |
| 10309 | g24 | 2014 | 38.0548 | -122.8332 | USA | CA | Drake 2 | 10.5600 | 2.5250 | detached | 13/15.1 | 13/15 |
| 10326 | g25 | 2014 | 38.0548 | -122.8332 | USA | CA | Drake 2 | 55.1934 | 2.9475 | closed bubble | 13/18 | 13/18 |
| 10327 | g25 | 2014 | 38.0548 | -122.8332 | USA | CA | Drake 2 | 48.4987 | 2.7791 | closed bubble | 13/18 | 13/18 |
| 10328 | g25 | 2014 | 38.0548 | -122.8332 | USA | CA | Drake 2 | 58.3688 | 3.1719 | closed bubble | 13/18 | 13/18 |
| 10329 | g25 | 2014 | 38.0548 | -122.8332 | USA | CA | Drake 2 | 9.4657 | 2.7828 | 3 | 13/18 | 13/18 |
| 10330 | g25 | 2014 | 38.0548 | -122.8332 | USA | CA | Drake 2 | 43.3505 | 2.7093 | closed bubble | 13/18 | 13/18 |
| 10331 | g25 | 2014 | 38.0548 | -122.8332 | USA | CA | Drake 2 | 50.3983 | 2.7992 | closed bubble | 13/18 | 13/18 |
| 10334 | g26 | 2014 | 38.0548 | -122.8332 | USA | CA | Drake 2 | 37.7795 | 2.7056 | closed bubble | 9/15.1 | 9/15 |
| 10347 | g19 | 2014 | 38.0552 | -122.8341 | USA | CA | Drake 3 | 44.2445 | 2.1129 | open bubble | 8/13 | 8/13 |

| | | | | | | | | | | | | |
|---|---|---|---|---|---|---|---|---|---|---|---|---|
| 10348 | g19 | 2014 | 38.0552 | -122.8341 | USA | CA | Drake 3 | 58.3031 | 2.3791 | open bubble | 8/13 | 8/13 |
| 10349 | g19 | 2014 | 38.0552 | -122.8341 | USA | CA | Drake 3 | 72.1097 | 2.6746 | complexed | 8/13 | 8/13 |
| 10350 | g19 | 2014 | 38.0552 | -122.8341 | USA | CA | Drake 3 | 74.8458 | 2.7096 | open bubble | 8/13 | 8/13 |
| 10354 | g19 | 2014 | 38.0552 | -122.8341 | USA | CA | Drake 3 | 26.0310 | 2.1869 | open bubble | 8/13 | 8/13 |
| 10355 | g19 | 2014 | 38.0552 | -122.8341 | USA | CA | Drake 3 | 68.2392 | 2.5192 | open bubble | 8/13 | 8/13 |
| 10356 | g19 | 2014 | 38.0552 | -122.8341 | USA | CA | Drake 3 | 84.6668 | 2.5712 | open bubble | 8/13 | 8/13 |
| 10380 | g19 | 2014 | 38.0552 | -122.8341 | USA | CA | Drake 3 | 82.1521 | 2.6304 | open bubble | 8/13 | 8/13 |
| 10384 | g19 | 2014 | 38.0552 | -122.8341 | USA | CA | Drake 3 | 65.6493 | 2.4653 | open bubble | 8/13 | 8/13 |
| 10502 | g27 | 2015 | 40.2125 | -8.4503 | Portugal | Coimbra | Agraria | 63.2363 | 3.0660 | closed bubble | 9/17 | 9/17 |
| 10503 | g28 | 2015 | 40.2125 | -8.4503 | Portugal | Coimbra | Agraria | 67.8892 | 2.9450 | closed bubble | 12.2/17 | 12.2/17 |
| 10504 | g29 | 2015 | 40.1222 | -8.2097 | Portugal | Lousã | Vilarinho | 83.8065 | 2.9968 | closed bubble | 7/14 | 7/14 |
| 10505 | g30 | 2015 | 40.1222 | -8.2097 | Portugal | Lousã | Vilarinho | 78.1388 | 2.9271 | closed bubble | 13/15.2 | 13/15 |
| 10506 | g31 | 2015 | 40.1222 | -8.2097 | Portugal | Lousã | Vilarinho | 63.0676 | 3.1471 | open bubble | 8/16.1 | 8/16 |
| 10508 | g31 | 2015 | 40.1222 | -8.2097 | Portugal | Lousã | Vilarinho | 60.6785 | 3.0980 | open bubble | 8/16.1 | 8/16 |
| 10509 | g31 | 2015 | 40.1222 | -8.2097 | Portugal | Lousã | Vilarinho | 69.2680 | 3.4496 | open bubble | 8/16.1 | 8/16 |
| 10510 | g32 | 2015 | 40.4575 | -8.7686 | Portugal | São Jacinto | Mira | 78.5979 | 2.7594 | closed bubble | 3/12.1 | 3/12.1 |
| 10511 | g33 | 2015 | 40.4575 | -8.7686 | Portugal | São Jacinto | Mira | 150.8590 | 3.0036 | complexed | 5/8 | 5/8 |
| 10512 | g33 | 2015 | 40.4575 | -8.7686 | Portugal | São Jacinto | Mira | 71.7597 | 3.4160 | complexed | 5/8 | 5/8 |
| 10513 | g34 | 2015 | 40.4575 | -8.7686 | Portugal | São Jacinto | Mira | 74.2309 | 3.1374 | closed bubble | 2/4 | 2/4 |
| 10707 | g15 | 2015 | 38.0547 | -122.8333 | USA | CA | Drake 2 | 66.6071 | 2.5081 | open bubble | 8/11 | 8/11 |
| 10708 | g15 | 2015 | 38.0547 | -122.8333 | USA | CA | Drake 2 | 75.6193 | 2.5813 | open bubble | 8/11 | 8/11 |
| 10709 | g13 | 2015 | 38.0547 | -122.8333 | USA | CA | Drake 2 | 69.3263 | 1.7702 | closed bubble | 11/- | 11/- |
| 10710 | g25 | 2015 | 38.0547 | -122.8333 | USA | CA | Drake 2 | 75.0543 | 3.2610 | closed bubble | 13/18 | 13/18 |
| 10711 | g25 | 2015 | 38.0547 | -122.8333 | USA | CA | Drake 2 | 61.4925 | 3.2394 | closed bubble | 13/18 | 13/18 |
| 10712 | g25 | 2015 | 38.0547 | -122.8333 | USA | CA | Drake 2 | 70.8210 | 3.1909 | closed bubble | 13/18 | 13/18 |
| 10713 | g24 | 2015 | 38.0547 | -122.8333 | USA | CA | Drake 2 | 70.4096 | 2.9695 | closed bubble | 13/15.1 | 13/15 |
| 10715 | g25 | 2015 | 38.0547 | -122.8333 | USA | CA | Drake 2 | 104.1500 | 3.0392 | closed bubble | 13/18 | 13/18 |
| 10716 | g35 | 2015 | 38.0547 | -122.8333 | USA | CA | Drake 2 | 55.9177 | 1.9615 | open bubble | 8/13 | 8/13 |
| 10717 | g36 | 2015 | 38.0545 | -122.8328 | USA | CA | Drake 2 | 61.8824 | 3.1861 | closed bubble | 11/13 | 11/13 |

| | | | | | | | | | | | | |
|---|---|---|---|---|---|---|---|---|---|---|---|---|
| 10718 | g37 | 2015 | 38.0545 | -122.8328 | USA | CA | Drake 2 | 48.7091 | 2.5721 | closed bubble | 13/15.2 | 13/15 |
| 10719 | g19 | 2015 | 38.0551 | -122.8342 | USA | CA | Drake 3 | 56.1117 | 2.3736 | open bubble | 8/13 | 8/13 |
| 10720 | g19 | 2015 | 38.0551 | -122.8342 | USA | CA | Drake 3 | 62.3315 | 2.5205 | open bubble | 8/13 | 8/13 |
| 10721 | g19 | 2015 | 38.0551 | -122.8342 | USA | CA | Drake 3 | 125.4650 | 2.3131 | open bubble | 8/13 | 8/13 |

* allele determination based on protein sequences. -: assembly incomplete

#### Supplementary Table 2. Specimens for rediscovering g21 and g22 homokaryotic individuals

| ID | Year | Latitude | Longitude | Country | Region | County | City | Site Name |
|---|---|---|---|---|---|---|---|---|
| 20009 | 2021 | 38.0552 | -122.8333 | USA | CA | Marin | Point Reyes National Seashore | Drake2 |
| 20010 | 2021 | 38.0550 | -122.8333 | USA | CA | Marin | Point Reyes National Seashore | Drake2 |
| 20011 | 2021 | 38.0550 | -122.8333 | USA | CA | Marin | Point Reyes National Seashore | Drake2 |
| 20012 | 2021 | 38.0550 | -122.8334 | USA | CA | Marin | Point Reyes National Seashore | Drake2 |
| 20013 | 2021 | 38.0546 | -122.8331 | USA | CA | Marin | Point Reyes National Seashore | Drake2 |
| 20014 | 2021 | 38.0547 | -122.8331 | USA | CA | Marin | Point Reyes National Seashore | Drake2 |
| 20015 | 2021 | 38.0546 | -122.8333 | USA | CA | Marin | Point Reyes National Seashore | Drake2 |
| 20016 | 2021 | 38.0548 | -122.8331 | USA | CA | Marin | Point Reyes National Seashore | Drake2 |
| 20017 | 2021 | 38.0547 | -122.8332 | USA | CA | Marin | Point Reyes National Seashore | Drake2 |
| 20018 | 2021 | 38.0545 | -122.8331 | USA | CA | Marin | Point Reyes National Seashore | Drake2 |
| 20030 | 2021 | 38.0547 | -122.8330 | USA | CA | Marin | Point Reyes National Seashore | Drake2 |
| 20032 | 2021 | 38.0550 | -122.8374 | USA | CA | Marin | Point Reyes National Seashore | Drake4 |
| 20033 | 2021 | 38.0550 | -122.8371 | USA | CA | Marin | Point Reyes National Seashore | Drake4 |
| 20034 | 2021 | 38.0552 | -122.8355 | USA | CA | Marin | Point Reyes National Seashore | Drake4 |
| 20036 | 2021 | 38.0547 | -122.8374 | USA | CA | Marin | Point Reyes National Seashore | Drake4 |
| 20041 | 2021 | 38.0546 | -122.8333 | USA | CA | Marin | Point Reyes National Seashore | Drake0 |
| 20043 | 2021 | 38.0544 | -122.8334 | USA | CA | Marin | Point Reyes National Seashore | Drake0 |
| 20046 | 2021 | 38.0553 | -122.8329 | USA | CA | Marin | Point Reyes National Seashore | Drake2 |
| 20047 | 2021 | 38.0552 | -122.9329 | USA | CA | Marin | Point Reyes National Seashore | Drake2 |
| 20048 | 2021 | 38.0553 | -122.8329 | USA | CA | Marin | Point Reyes National Seashore | Drake2 |
| 20049 | 2021 | 38.0554 | -122.8329 | USA | CA | Marin | Point Reyes National Seashore | Drake2 |
| 20050 | 2021 | 38.0549 | -122.8330 | USA | CA | Marin | Point Reyes National Seashore | Drake2 |
| 20051 | 2021 | 38.0548 | -122.8330 | USA | CA | Marin | Point Reyes National Seashore | Drake2 |
| 20052 | 2021 | 38.0546 | -122.8327 | USA | CA | Marin | Point Reyes National Seashore | Drake2 |
| 20053 | 2021 | 38.0546 | -122.8328 | USA | CA | Marin | Point Reyes National Seashore | Drake2 |
| 20054 | 2021 | 38.0545 | -122.8330 | USA | CA | Marin | Point Reyes National Seashore | Drake2 |

| | | | | | | | | |
|---|---|---|---|---|---|---|---|---|
| 20055 | 2021 | 38.0546 | -122.8326 | USA | CA | Marin | Point Reyes National Seashore | Drake2 |
| 20056 | 2021 | 38.0546 | -122.8326 | USA | CA | Marin | Point Reyes National Seashore | Drake2 |
| 20058 | 2021 | 38.0547 | -122.8325 | USA | CA | Marin | Point Reyes National Seashore | Drake2 |
| 20060 | 2021 | 38.0547 | -122.8323 | USA | CA | Marin | Point Reyes National Seashore | Drake2 |

#### Supplementary Table 3. Primers used for detecting heterozygosity in specimens without whole genome sequencing data

| Primer ID | Forward | Reverse | Amplicon Length | Contig | Start | End | SNP sites | Functional Annotation | Nick Name | P(hom) (CA)* | P(hom) (All)* |
|---|---|---|---|---|---|---|---|---|---|---|---|
| 00g060260-3 | ACC GCC AGA CTT GTC AAA CA | TCA ATC GCA GCA GTG GAA CT | 691 | Contig87 | 400939 | 401629 | multiple | beta-flanking gene | | 0.13† | 0.15† |
| 00g060260-16 | CAG CCT GGA CAA TCT CGT CA | TCA ATC GCA GCA GTG GAA CT | 248 | Contig87 | 401382 | 401629 | multiple | beta-flanking gene | | 0.31† | 0.37† |
| 00g000510-2 | TTC AGC CGA CGT TAC GAC TC | TGG GAG TTG GCG TTG TTT CT | 220 | Contig0 | 327946 | 328165 | 328045A/G | Eukaryotic translation initiation factor 3 subunit C | Het-1 | 0.75 | 0.57 |
| 00g004840-9 | AAG CCT AAA GGT GGC GTT CA | GCG AGA TCG ACC GCA TAG TT | 137 | Contig2 | 766786 | 766922 | 766833G/A; 766848C/T; 766902T/C | Fimbrin | Het-2 | 0.58 | 0.43 |
| 00g009240-3 | GCT ATT GTC CAG GAG GTC CA | GCC TAG ACT TTT CAG CTG CTC | 149 | Contig5 | 299743 | 299891 | 299822A/C; 299864G/A | Elongator complex protein 3 | Het-3 | 0.55 | 0.51 |
| 00g011220-3 | GGA TAT GCG GAC CAG GAT CG | TCG TCC TCG AAG TAA GCA TCC | 153 | Contig5 | 748470 | 748622 | 748593G/C | Transcription elongation factor Spt6 | Het-4 | 0.31 | 0.29 |
| 00g016590-6 | CAC TTT GCG CAG ACT CAC G | GTA AAC AGT CCG AAG GGG CT | 172 | Contig7 | 692570 | 692741 | 692691T/C | RNA polymerase I-specific transcription initiation factor RRN3 | Het-5 | 0.53 | 0.52 |
| 00g021750-9 | AGA TCC CGA AGT TCA CTT CAA A | AGT TCT TGA CCT TCT CTG GGT | 116 | Contig8 | 62025 | 62140 | 62082T/C | Clathrin heavy chain | Het-6 | 0.53 | 0.54 |
| 00g023990-5 | GCA TCT TCG TAC GGG ACA GT | GCA GCT TGG CTT TCG TCA AT | 156 | Contig9 | 207878 | 208033 | 207956A/G; 208004A/G | DNA-directed RNA polymerase | Het-7 | 0.51 | 0.49 |
| 00g053490-1 | ATC TCA CAC AGC CAG GTT GG | TAG GGA TGA CCC TCC TTC GG | 133 | Contig63 | 110061 | 110193 | 110102G/A; 110119T/C | DNA polymerase gamma | Het-9 | 0.52 | 0.50 |
| 00g072790-8 | TCC AAG AAC AAT GAC CGG GAA | ATC TCG GAG TCG GTT CCT TTG | 143 | Contig104 | 676464 | 676606 | 676503G/A; 676536G/A | GPI-anchored wall transfer protein | Het-10 | 0.47 | 0.43 |
| 00g083180-4 | AAC GAA GTG GAA TGC GCG A | TAG GCC GCA TCA CAC AAG AC | 111 | Contig166 | 99533 | 99643 | 99599T/G | DNA polymerase alpha subunit B | Het-11 | 0.51 | 0.51 |

* Estimates of probability of homozygosity are from clone corrected dataset, without haploids, of either Californian samples or all samples

† Due to the complexity of these two regions, the probability of homozygosity is estimated with clipping off 80 nucleotides of either ends of the amplicon, others are estimated by the exact observable sites in Sanger sequences

#### Supplementary Table 4. Plasmids used for yeast two-hybrid assays

| Plasmid | Description | Reference |
|---|---|---|
| pCH478 (pGAD-C1) | pGAD-C1 | James et al., 1996 [61] |
| pCH312 (pGBD-C1) | pGBD-C1 | James et al., 1996 [61] |
| pGAD-C1-HD1-5 | HD1 ORF of allele 5 fused to the *GAL4* activation domain | This study |
| pGAD-C1-HD1-8 | HD1 ORF of allele 8 fused to the *GAL4* activation domain | This study |
| pGAD-C1-HD1-13 | HD1 ORF of allele 13 fused to the *GAL4* activation domain | This study |
| pGAD-C1-HD2-5 | HD2 ORF of allele 5 fused to the *GAL4* activation domain | This study |
| pGAD-C1-HD2-8 | HD2 ORF of allele 8 fused to the *GAL4* activation domain | This study |
| pGAD-C1-HD2-13 | HD2 ORF of allele 13 fused to the *GAL4* activation domain | This study |
| pGBD-C1-HD1-5 | HD1 ORF of allele 5 fused to the *GAL4* DNA binding domain | This study |
| pGBD-C1-HD1-8 | HD1 ORF of allele 8 fused to the *GAL4* DNA binding domain | This study |
| pGBD-C1-HD1-13 | HD1 ORF of allele 13 fused to the *GAL4* DNA binding domain | This study |
| pGBD-C1-HD2-5 | HD2 ORF of allele 5 fused to the *GAL4* DNA binding domain | This study |
| pGBD-C1-HD2-8 | HD2 ORF of allele 8 fused to the *GAL4* DNA binding domain | This study |
| pGBD-C1-HD2-13 | HD2 ORF of allele 13 fused to the *GAL4* DNA binding domain | This study |

#### Supplementary Table 5. Yeast strain used for yeast two-hybrid assays

| Strain | Genotype | Reference |
|---|---|---|
| CHY1268 (PJ69-4a) | MATa trp1-901 leu2-3, 112 ura3-52 his3-200 gal4*D gal80D LYS::GAL1-HIS3 GAL2-ADE2 met2::GAL7-lacZ* | James et al., 1996 [91] |

**Supplementary Table 6.** Additional specimens without whole genome sequencing data screened as potentially homokaryotic sporocarps

| ID | Year | Latitude | Longitude | Country | Region | County | City | Site Name | Determiner | Collector |
|---|---|---|---|---|---|---|---|---|---|---|
| 10001 | 2006 | NA | NA | UK | NA | NA | Middlesex | Hampstead Heath Extension | Overall | Overall |
| 10009 | 2004 | NA | NA | UK | North Hampshire | NA | Shortheath Common | near Bordon | Legon | Legon |
| 10012 | 1999 | NA | NA | UK | West Sussex | NA | Petworth | The Mens and the Cut Nat. Reserve | Legon | Legon |
| 10013 | 2004 | NA | NA | UK | Surrey | NA | Sheepleas | Green Dene (near West Horsley) | Legon | Legon |
| 10015 | 2001 | NA | NA | UK | Surrey | NA | NA | Kew Royal Botanical Gardens | Brown | Brown |
| 10017 | 2004 | NA | NA | UK | West Sussex | NA | NA | The Marlows | Legon | Legon |
| 10021 | 2005 | NA | NA | UK | West Suffolk | NA | Mildenhall | Mildenhall Woods | Legon | Legon |
| 10022 | 2004 | NA | NA | UK | South Essex | NA | NA | Epping Forest near Earl's Path | Overall | Overall |
| 10024 | 1994 | NA | NA | UK | Surrey | NA | Mickelham | Norbury Park | Legon | Legon |
| 10025 | 1996 | NA | NA | UK | Oxfordshire | NA | Bix | The Warburg Preserve | Legon | Legon |
| 10026 | 1997 | NA | NA | UK | Oxfordshire | NA | Bix | The Warburg Preserve | Legon | Legon |
| 10027 | 1997 | NA | NA | UK | South Somerset | NA | Staple Common | near Castle Neroche | Legon | Legon |
| 10028 | 1998 | NA | NA | UK | Surrey | NA | NA | Kew Royal Botanical Gardens | Legon | Legon |
| 10029 | 1998 | NA | NA | UK | North Somerset | NA | NA | Stree in Great Breech Woods | Legon | Legon |
| 10031 | 2006 | 39.1899 | -74.8535 | USA | NJ | Cape May | Dennisville | Jake's Landing Rd | Wolfe | Wolfe |
| 10034 | 2006 | 39.1899 | -74.8535 | USA | NJ | Cape May | Dennisville | Jake's Landing Rd | Wolfe | Wolfe |
| 10038 | 2006 | 39.1899 | -74.8535 | USA | NJ | Cape May | Dennisville | Jake's Landing Rd | Wolfe | Wolfe |
| 10040 | 2006 | 39.1899 | -74.8535 | USA | NJ | Cape May | Dennisville | Jake's Landing Rd | Wolfe | Wolfe |
| 10044 | 2006 | 39.1899 | -74.8535 | USA | NJ | Cape May | Dennisville | Jake's Landing Rd | Wolfe | Wolfe |
| 10048 | 2006 | 39.1899 | -74.8535 | USA | NJ | Cape May | Dennisville | Jake's Landing Rd | Wolfe | Wolfe |
| 10051 | 2006 | 39.1899 | -74.8535 | USA | NJ | Cape May | Dennisville | Jake's Landing Rd | Wolfe | Wolfe |
| 10053 | 2006 | 39.1899 | -74.8535 | USA | NJ | Cape May | Dennisville | Jake's Landing Rd | Wolfe | Wolfe |
| 10054 | 2006 | 39.1899 | -74.8535 | USA | NJ | Cape May | Dennisville | Jake's Landing Rd | Wolfe | Wolfe |
| 10056 | 2006 | 39.1899 | -74.8535 | USA | NJ | Cape May | Dennisville | Jake's Landing Rd | Wolfe | Wolfe |
| 10061 | 2006 | 39.1899 | -74.8535 | USA | NJ | Cape May | Dennisville | Jake's Landing Rd | Wolfe | Wolfe |
| 10067 | 2006 | 39.1899 | -74.8535 | USA | NJ | Cape May | Dennisville | Jake's Landing Rd | Wolfe | Wolfe |

| | | | | | | | | | | |
|---|---|---|---|---|---|---|---|---|---|---|
| 10069 | 2006 | 39.1899 | -74.8535 | USA | NJ | Cape May | Dennisville | Jake's Landing Rd | Wolfe | Wolfe |
| 10072 | 2006 | 39.1899 | -74.8535 | USA | NJ | Cape May | Dennisville | Jake's Landing Rd | Wolfe | Wolfe |
| 10076 | 2006 | 39.1899 | -74.8535 | USA | NJ | Cape May | Dennisville | Jake's Landing Rd | Wolfe | Wolfe |
| 10109 | 2007 | NA | NA | USA | NY | Monroe | Rochester | Durand Eastman Park | Wolfe, Richard | Wolfe, Richard |
| 10110 | 2007 | NA | NA | USA | NY | Monroe | Rochester | Durand Eastman Park | Wolfe, Richard | Wolfe, Richard |
| 10112 | 2007 | NA | NA | USA | NY | Monroe | Rochester | Durand Eastman Park | Wolfe, Richard | Wolfe, Richard |
| 10113 | 2007 | NA | NA | USA | NY | Monroe | Rochester | Durand Eastman Park | Wolfe, Richard | Wolfe, Richard |
| 10114 | 2007 | NA | NA | USA | NY | Monroe | Rochester | Durand Eastman Park | Wolfe, Richard | Wolfe, Richard |
| 10124 | 2007 | NA | NA | USA | NY | Monroe | Rochester | Durand Eastman Park | Wolfe, Richard | Wolfe, Richard |
| 10128 | 2007 | NA | NA | USA | NY | Monroe | Rochester | Durand Eastman Park | Wolfe, Richard | Wolfe, Richard |
| 10131 | 2007 | NA | NA | USA | NY | Monroe | Rochester | Durand Eastman Park | Wolfe, Richard | Wolfe, Richard |
| 10132 | 2007 | NA | NA | USA | NY | Monroe | Rochester | Durand Eastman Park | Wolfe, Richard | Wolfe, Richard |
| 10134 | 2007 | NA | NA | USA | NY | Monroe | Rochester | Durand Eastman Park | Wolfe, Richard | Wolfe, Richard |
| 10141 | 2001 | 37.8949 | -122.3047 | USA | CA | Alameda | Albany | Albany Hill | Bruns | Bruns |
| 10142 | 2001 | 37.8949 | -122.3047 | USA | CA | Alameda | Albany | Albany Hill | Bruns | Bruns |
| 10151 | 2001 | 37.8949 | -122.3047 | USA | CA | Alameda | Albany | Albany Hill | Bruns | Bruns |
| 10153 | 2001 | 37.8949 | -122.3047 | USA | CA | Alameda | Albany | Albany Hill | Bruns | Bruns |
| 10157 | 2001 | 37.8949 | -122.3047 | USA | CA | Alameda | Albany | Albany Hill | Bruns | Bruns |
| 10158 | 2001 | 37.8949 | -122.3047 | USA | CA | Alameda | Albany | Albany Hill | Bruns | Bruns |
| 10159 | 2001 | 37.8949 | -122.3047 | USA | CA | Alameda | Albany | Albany Hill | Bruns | Bruns |
| 10162 | 2001 | 37.7769 | -122.1470 | USA | CA | Alameda | Oakland | Leona Canyon | Bruns | Pringle |
| 10166 | 2001 | 37.7769 | -122.1470 | USA | CA | Alameda | Oakland | Leona Canyon | Bruns | Pringle |
| 10167 | 2001 | 37.7769 | -122.1470 | USA | CA | Alameda | Oakland | Leona Canyon | Bruns | Pringle |
| 10192 | 2002 | NA | NA | Canada | British Colombia | Victoria Island | Victoria | Budget Rent a Car/Crystal Gardens | McCann | McCann |
| 10193 | 2002 | NA | NA | Canada | British Colombia | Victoria Island | Victoria | NA | Dennis | Dennis |
| 10194 | 2000 | NA | NA | Canada | British Colombia | Victoria Island | Victoria | NA | Dennis | Dennis |
| 10195 | 2002 | NA | NA | Canada | British Colombia | Victoria Island | Victoria | Downtown in garden | Dennis | Dennis |
| 10196 | 2002 | NA | NA | Canada | British Colombia | Victoria Island | Victoria | NA | Dennis | Dennis |
| 10197 | 1998 | NA | NA | Canada | British Colombia | Victoria Island | Victoria | Rockland Ave | Callan | Fleck |

| | | | | | | | | | | |
|---|---|---|---|---|---|---|---|---|---|---|
| 10198 | 1998 | NA | NA | Canada | British Colombia | Victoria Island | Victoria | Rockland Ave | Callan, Windler | Howe |
| 10199 | 1997 | NA | NA | Canada | British Colombia | NA | Mission | NA | Kroeger | Tatum |
| 10270 | 2002 | NA | NA | USA | CA | Alameda | Berkeley | Tilden Park | NA | Pringle |
| 10271 | 2002 | NA | NA | USA | CA | Alameda | Berkeley | Tilden Park | NA | Pringle |
| 10272 | 2002 | NA | NA | USA | CA | Alameda | Berkeley | Tilden Park | NA | Pringle |
| 10273 | 2002 | NA | NA | USA | CA | Alameda | Berkeley | Tilden Park | NA | Pringle |
| 10274 | 2002 | NA | NA | USA | CA | Alameda | Berkeley | Tilden Park | NA | Pringle |
| 11002 | 2019 | NA | NA | France | Occitanie | NA | Montpellier | NA | Franck | Franck |
| 11003 | 2019 | NA | NA | France | Occitanie | NA | Montpellier | NA | Franck | Franck |
| 11007 | 2019 | NA | NA | France | Occitanie | NA | Montpellier | NA | Franck | Franck |
| 11012 | 2019 | NA | NA | France | Occitanie | NA | Montpellier | NA | Franck | Franck |
| 11016 | 2019 | NA | NA | France | Occitanie | NA | Montpellier | NA | Franck | Franck |
| 11020 | 2019 | NA | NA | France | Occitanie | NA | Montpellier | NA | Franck | Franck |
| 11022 | 2019 | NA | NA | France | Occitanie | NA | Montpellier | NA | Franck | Franck |
| 11023 | 2019 | NA | NA | France | Occitanie | NA | Montpellier | NA | Franck | Franck |
| 11024 | 2019 | NA | NA | France | Occitanie | NA | Montpellier | NA | Franck | Franck |
| 11029 | 2019 | NA | NA | France | Occitanie | NA | Montpellier | NA | Franck | Franck |
| 11031 | 2019 | NA | NA | France | Occitanie | NA | Montpellier | NA | Franck | Franck |
| 11033 | 2019 | NA | NA | France | Occitanie | NA | Montpellier | NA | Franck | Franck |
| 11051 | 2019 | NA | NA | Norway | Hordaland | NA | Bomlo | Aphal1-Stord | Wollan | Wollan |
| 11052 | 2019 | NA | NA | Norway | Hordaland | NA | Bomlo | Aphal1-Stord | Wollan | Wollan |
| 11053 | 2019 | NA | NA | Norway | Hordaland | NA | Tysnet | Aphal2-Stord | Wollan | Wollan |
| 11054 | 2019 | NA | NA | Norway | Hordaland | NA | Tysnet | Aphal2-Stord | Wollan | Wollan |
| 11055 | 2019 | NA | NA | Norway | Hordaland | NA | Tysnet | Aphal2-Stord | Wollan | Wollan |
| 11056 | 2019 | NA | NA | Norway | Hordaland | NA | Tysnet | Aphal2-Stord | Wollan | Wollan |
| 11101 | 2019 | 48.2472 | 16.2458 | Austria | Niederösterreich | NA | Tulln | Exelberg | Krisai-Greilhuber | Krisai-Greilhuber |
| 11102 | 2019 | 48.2472 | 16.2458 | Austria | Niederösterreich | NA | Tulln | Exelberg | Krisai-Greilhuber | Krisai-Greilhuber |
| 11103 | 2019 | 48.2472 | 16.2458 | Austria | Niederösterreich | NA | Tulln | Exelberg | Krisai-Greilhuber | Krisai-Greilhuber |
| 11104 | 2019 | 48.1533 | 16.2475 | Austria | Liesing | NA | Wien | Maurerwald | Krisai-Greilhuber | Krisai-Greilhuber |

| | | | | | | | | | | |
|---|---|---|---|---|---|---|---|---|---|---|
| 11105 | 2019 | 48.1533 | 16.2475 | Austria | Liesing | NA | Wien | Maurerwald | Krisai-Greilhuber | Krisai-Greilhuber |
| 11106 | 2019 | 48.1533 | 16.2475 | Austria | Liesing | NA | Wien | Maurerwald | Krisai-Greilhuber | Krisai-Greilhuber |
| 11107 | 2019 | 48.1533 | 16.2475 | Austria | Liesing | NA | Wien | Maurerwald | Krisai-Greilhuber | Krisai-Greilhuber |
| 11108 | 2019 | 48.1533 | 16.2475 | Austria | Liesing | NA | Wien | Maurerwald | Krisai-Greilhuber | Krisai-Greilhuber |
| 11109 | 2019 | 48.1533 | 16.2475 | Austria | Liesing | NA | Wien | Maurerwald | Krisai-Greilhuber | Krisai-Greilhuber |
| 11110 | 2019 | 48.1533 | 16.2475 | Austria | Liesing | NA | Wien | Maurerwald | Krisai-Greilhuber | Krisai-Greilhuber |
| 11111 | 2019 | 48.1533 | 16.2475 | Austria | Liesing | NA | Wien | Maurerwald | Krisai-Greilhuber | Krisai-Greilhuber |
| 11112 | 2019 | 42.5667 | 16.5606 | Austria | Burgenland | NA | Horitschon | Ragerwald | Krisai-Greilhuber | Krisai-Greilhuber |
| 11113 | 2019 | 48.2231 | 16.2567 | Austria | Hernals | NA | Wien | Moosgraben | Gross | Gross |
| 11121 | 2017 | 58.6053 | 23.6981 | Estonia | NA | Pärnu | Karuse | NA | Zettur | Zettur |
| 11122 | 2016 | 58.9653 | 22.5238 | Estonia | NA | Hiiu | Pihla | NA | Saar | Ploompuu |
| 11131 | 2019 | 46.1801 | 20.1368 | Hungary | Hungary | Csongrád | Csongrád | Tisza | Harrow | Harrow |
| 11132 | 2019 | 46.1801 | 20.1368 | Hungary | Hungary | Csongrád | Csongrád | Tisza | Harrow | Harrow |
| 11133 | 2019 | 46.1801 | 20.1368 | Hungary | Hungary | Csongrád | Csongrád | Tisza | Harrow | Harrow |
| 11134 | 2019 | 46.4372 | 20.1201 | Hungary | Hungary | Csongrád | Csongrád | Doc | Harrow | Harrow |
| 11135 | 2019 | 46.4372 | 20.1201 | Hungary | Hungary | Csongrád | Csongrád | Doc | Harrow | Harrow |
| 11137 | 2019 | 46.4372 | 20.1201 | Hungary | Hungary | Csongrád | Csongrád | Doc | Harrow | Harrow |
| 11138 | 2019 | 46.4372 | 20.1201 | Hungary | Hungary | Csongrád | Csongrád | Doc | Harrow | Harrow |
| 11141 | 2019 | 46.4372 | 20.1201 | Hungary | Hungary | Csongrád | Csongrád | Doc | Harrow | Harrow |
| 11142 | 2019 | 46.4372 | 20.1201 | Hungary | Hungary | Csongrád | Csongrád | Doc | Harrow | Harrow |
| 11143 | 2019 | 46.4372 | 20.1201 | Hungary | Hungary | Csongrád | Csongrád | Doc | Harrow | Harrow |
| 11144 | 2019 | 46.4372 | 20.1201 | Hungary | Hungary | Csongrád | Csongrád | Doc | Harrow | Harrow |
| 11145 | 2019 | 46.4372 | 20.1201 | Hungary | Hungary | Csongrád | Csongrád | Doc | Harrow | Harrow |
| 11151 | 2019 | 47.3629 | 8.4844 | Switzerland | Switzerland | Zürich | Zürich | Birm | Harrow | Harrow |
| 11152 | 2019 | 47.3629 | 8.4844 | Switzerland | Switzerland | Zürich | Zürich | Birm | Harrow | Harrow |

#### Supplementary Table 7. SNP genotypes of known and putative homokaryotic specimens

| SNP sites | Known | | Putative* | | | | | | | | |
|---|---|---|---|---|---|---|---|---|---|---|---|
| | 10224 | 10293 | 10028 | 10038 | 10044 | 10048 | 10051 | 11108 | 20018 | 20030 | 20034 |
| Contig0:328045A/G | G | G | A | A | A | A | A | A | G | G | G |
| Contig2:766833G/A | A | G | - | - | - | - | - | - | - | - | - |
| Contig2:766848C/T | T | C | T | C | C/T | C/T | C | C | C | T | T |
| Contig2:766902T/C | T | T | T | T | T | T | T | T | T | T | T |
| Contig5:299822A/C | A | A | A | A | A | A | A | A | A | A | A |
| Contig5:299864G/A | A | A | A/G | A | A/G | A/G | A/G | A/G | A | A | A |
| Contig5:748593G/C | C | C | G/C | G/C | G/C | G | G | G/C | C | C | C |
| Contig7:692691T/C | T | C | C | C | C | C | C | C | C | T | T |
| Contig8:62082T/C | T | T | C | C | C | C/T | C | C | T | T | T |
| Contig9:207956A/G | G | A | A | A | A/G | A/G | A | A/G | A | G | G |
| Contig9:208004A/G | G | A | A | A | A/G | A/G | A | A/G | A | G | G |
| Contig63:110102G/A | G | A | - | - | - | - | - | - | - | - | - |
| Contig63:110119T/C | T | C | C | C | C | C/T | T | C | C | T | T |
| Contig104:676503G/A | G | G | - | - | - | - | - | - | - | G | - |
| Contig104:676536G/A | G | G | A | A | A | A/G | A/G | A | G | G | G |
| Contig166:99599T/G | G | T | T | T/G | T | T | T | T/G | T | G | G |

* Each specimen possesses a homozygous beta-flanking gene
